# EHMT1 and EHMT2 Control the Timing of Cortical Neurogenesis Through H3K9me2-Dependent and -Independent Mechanisms

**DOI:** 10.64898/2026.11.16.767207

**Authors:** Adhyeta Choudhuri, Chris Hemmerich, Ram Podicheti, Sungyun Kang, José-Manuel Baizabal

## Abstract

The precise timing of neural progenitor differentiation and cell fate specification is critical for establishing the size and cellular diversity of the cerebral cortex. However, how chromatin regulators control the temporal progression of neurogenesis remains poorly understood. Here we investigate the cooperative roles of Euchromatic Histone Lysine Methyltransferases 1 and 2 (EHMT1 and EHMT2) in mouse cortical neurogenesis. We demonstrate that EHMT1– and EHMT2-mediated H3K9me2 deposition is dispensable for H3K9me3, and that these histone marks occupy distinct chromatin regions. Combined loss of EHMT1 and EHMT2 reduces neural progenitor proliferation, promotes differentiation, and disrupts the transition from deep– to upper-layer neurogenesis, resulting in microcephaly and altered neuronal subtype composition in the postnatal cortex. We further show that EHMT2 directly regulates genes involved in neurogenesis and upper-layer neuron specification through both transcriptional repression and activation. Moreover, EHMT1 and EHMT2 suppress the premature expression of synaptic genes in neural progenitors through complementary mechanisms involving H3K9me2-associated chromatin compaction and H3K9me2-independent silencing in cooperation with PRDM16. Together, these findings establish EHMT1 and EHMT2 as central regulators of the temporal control of cortical neurogenesis and reveal molecular mechanisms that may contribute to the pathogenesis of Kleefstra syndrome.

## INTRODUCTION

The progression of cortical neurogenesis depends on the precisely coordinated differentiation of neural stem cells (NSCs, also known as radial glia) into intermediate progenitor cells (IPCs) and, ultimately, cortical neurons (1). During early neurogenesis, NSCs and IPCs, collectively referred to as neural progenitors, predominantly produce deep-layer neurons (1). As development proceeds, a temporal shift in neuronal fate initiates the production of upper-layer neurons (1). Premature neuronal differentiation can deplete the progenitor pool, reducing the overall neuronal output and potentially causing microcephaly (2). Thus, the proper temporal control of cortical neurogenesis requires mechanisms that transiently restrain neuronal differentiation, preserving the proliferative progenitor population for continued cortical expansion.

Histone methylation is associated with cell-type-specific gene expression programs and is an important regulator of cell fate specification and differentiation (3). Histone H3 lysine 9 (H3K9) can undergo three distinct methylation states, each potentially contributing differently to transcriptional regulation (4). H3K9 monomethylation (H3K9me1) is associated with active genes, whereas di– and trimethylation (H3K9me2 and H3K9me3) are linked to transcriptional repression (5). H3K9me2 is enriched within lamina-associated domains (LADs), where it is thought to contribute to the silencing of genes and transposable elements at the nuclear periphery (6–8). This role suggests that H3K9me2 is sufficient to promote gene silencing, rather than serving solely as a transitional state to H3K9me3-mediated repression (9). Moreover, H3K9me2 and H3K9me3 mark common and distinct genomic regions in cell lines, further suggesting that these histone modifications have redundant and unique regulatory roles (5).

H3K9me1 and H3K9me2 are deposited by Euchromatic Histone Lysine Methyltransferase 1 (EHMT1, also known as GLP) and EHMT2 (also known as G9a) (4). Both enzymes contain a carboxy-terminal SET (Su(var)3-9, Enhancer-of-zeste, and Trithorax) domain that confers methyltransferase activity (4). EHMT1 and EHMT2 (hereafter EHMT1/2) form homo– and heterodimeric complexes, with the heterodimer exhibiting the more robust catalytic activity and pronounced effects on gene regulation (10–12). *Ehmt1/2* are ubiquitously expressed, and individual loss of either *Ehmt1* or *Ehmt2* in mouse embryos causes similar developmental abnormalities and early embryonic lethality, consistent with their cooperative functions (10, 11). In addition, the combined loss of EHMT1/2 results in the depletion of H3K9me1 and H3K9me2 and the concomitant derepression of gene expression in early mouse embryos and embryonic stem cells (10, 11).

In the postnatal mouse brain, neuronal deletion of *Ehmt1/2* leads to the derepression of a subset of non-neuronal genes and is accompanied by behavioral deficits (13). EHMT1/2 also contribute to cortical synaptic homeostasis through transcriptional repression (14). In olfactory neurons, EHMT1/2 are required for the proper silencing of olfactory receptor genes, thereby promoting olfactory neuron diversification in mice (15). In humans, *EHMT1* haploinsufficiency causes Kleefstra syndrome, a neurodevelopmental disorder characterized by intellectual disability, microcephaly, and autistic features (16, 17). In addition, emerging evidence implicates *EHMT2* dysfunction in Kleefstra-like neurodevelopmental phenotypes (18).

Despite the established roles of EHMT1/2 in neuronal function and Kleefstra syndrome, their cooperative functions in cortical development remain poorly understood. To address this gap, we generated cortex-specific *Ehmt1/2* double-knockout (DKO) mice at the onset of embryonic neurogenesis. Our findings reveal that EHMT1/2 are essential for sustaining neural progenitor proliferation and regulating the temporal progression from deep– to upper-layer neuron production. We further demonstrate that EHMT1/2 regulate genes associated with neurogenesis and upper-layer neuron specification. More broadly, EHMT1/2 exert a dual regulatory role by primarily repressing synaptic genes while promoting the expression of metabolic genes. Notably, these regulatory activities are only partially dependent on H3K9me2, indicating that EHMT1/2 control cortical neurogenesis through mechanisms extending beyond their canonical histone methyltransferase activity. Consistent with this, we identify a cooperative interaction between EHMT1/2 and PRDM16 that promotes the silencing of synaptic genes in neural progenitors largely independently of H3K9me2-marked chromatin. Collectively, our findings uncover distinct transcriptional and chromatin-associated functions of EHMT1/2 that coordinate gene expression programs in neural progenitors to ensure proper cortical growth and neuronal subtype composition.

## RESULTS

### EHMT1/2 specifically generate H3K9me2-marked chromatin in the embryonic cortex

To investigate the roles of EHMT1/2 in the embryonic cortex, we generated cortex-specific loss-of-function mutants before the onset of cortical neurogenesis at embryonic day 9.5 (E9.5) by crossing *Emx1-Cre* mice with mice carrying conditional floxed alleles of *Ehmt1* and *Ehmt2* (13, 19, 20). This strategy yielded single-knockout (SKO) mice for each gene and DKO mice. Immunostaining confirmed a substantial reduction in EHMT1/2 levels in the E15.5 DKO cortex (Fig. 1A). H3K9me2 levels were markedly reduced in neural progenitors of the E15.5 *Ehmt1* SKO cortex and moderately reduced in the *Ehmt2* SKO cortex compared with wild-type (WT) floxed controls (Fig. 1B). The near-complete depletion of H3K9me2 in the *Ehmt1* SKO cortex was accompanied by a pronounced reduction in EHMT2 levels (Fig. S1A), consistent with previous findings that EHMT1 regulates EHMT2 stability (11). In addition, H3K9me2 depletion was slightly more pronounced in the DKO cortex than in the *Ehmt1* SKO cortex (Fig. 1B). Loss of H3K9me2 persisted in neurons of the postnatal day 10 (P10) DKO cortex (Fig. S1B). These findings indicate that EHMT1/2 cooperate to establish and maintain H3K9me2 in the developing cortex.

**Figure 1.**
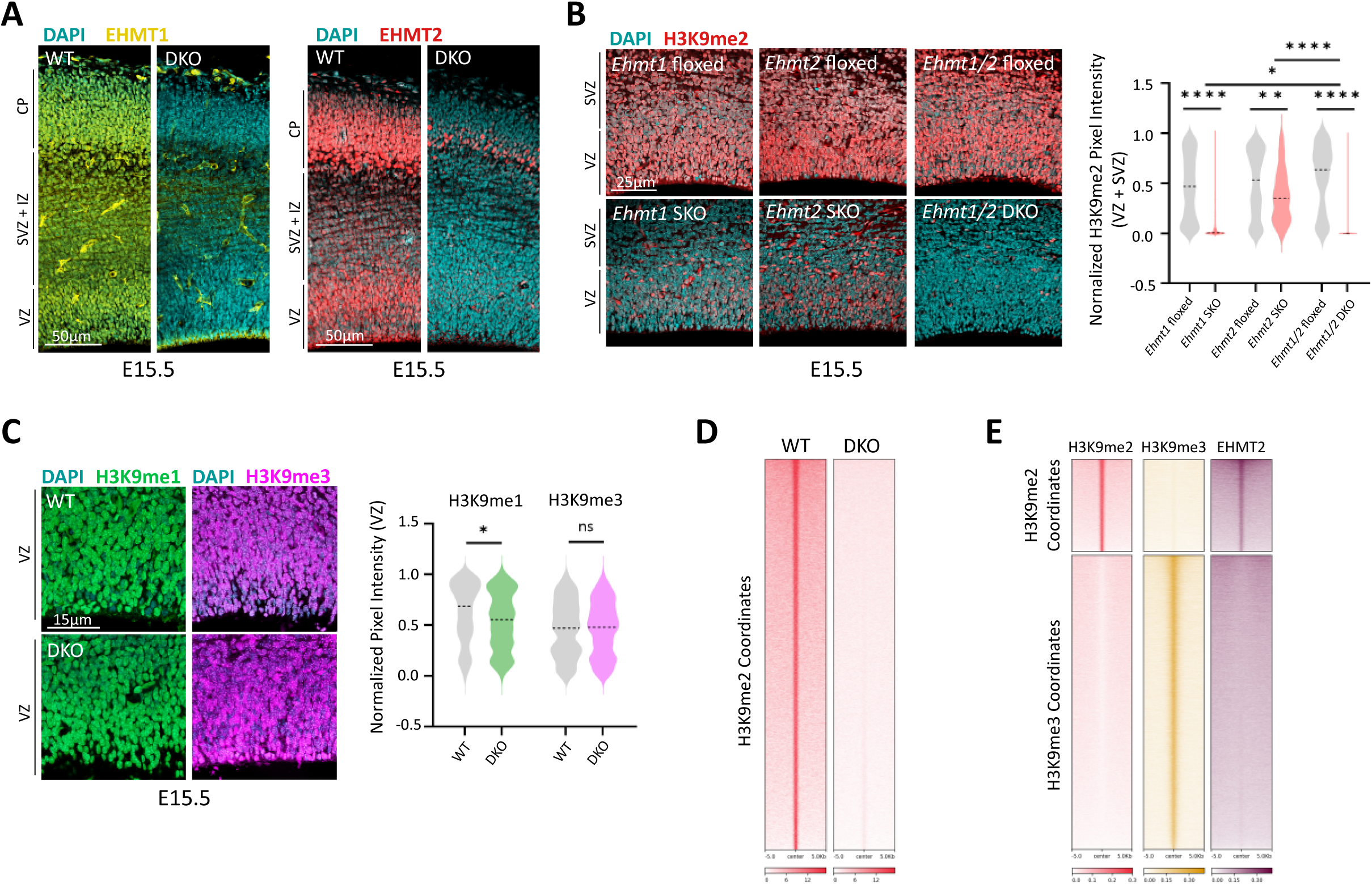
EHMT1 and EHMT2 cooperate to establish H3K9me2 at genomic regions distinct from H3K9me3. **(A)** Immunostaining showing ubiquitous EHMT1 and EHMT2 detection in the WT E15.5 cortex and their loss in the DKO cortex. **(B)** Immunostaining of E15.5 cortices showing robust H3K9me2 depletion in the VZ and SVZ of the *Ehmt1* SKO, moderate depletion in the *Ehmt2* SKO, and near-complete depletion in the *Ehmt1/2* DKO compared with WT floxed controls (n = 3). **(C)** Immunostaining of E15.5 cortices showing partial loss of H3K9me1 and no significant change in H3K9me3 in the DKO cortex (n = 3). **(D)** ChIP-seq heatmaps showing spike-in-normalized H3K9me2 enrichment in E14.5 WT cortices and the near-complete loss of H3K9me2 in DKO cortices. **(E)** ChIP-seq heatmaps showing that H3K9me2 and H3K9me3 occupy distinct chromatin regions in the E14.5 cortex. EHMT2-binding only overlaps with H3K9me2. Heatmaps were sorted by mean signal intensity across all samples, with the highest signal at the top. VZ, ventricular zone; SVZ, subventricular zone; IZ, intermediate zone; CP, cortical plate. Data are presented as mean ± SEM. Statistical significance was determined using a two-tailed unpaired *t*-test; ns, not significant.

We next examined whether EHMT1/2 loss affected other H3K9 methylation states. Unexpectedly, H3K9me1 levels were only marginally reduced in the E15.5 DKO cortex (Fig. 1C), contrasting with previous findings that EHMT1/2 generate most H3K9me1 in early mouse embryos and embryonic stem cells (10, 11). Moreover, H3K9me3 levels remained unchanged in the E15.5 DKO cortex (Fig. 1C). Consistent with these findings, H3K9me1 and H3K9me3 levels were also comparable between WT and DKO cortices at P10 (Fig. S1B). These results indicate that H3K9me2 is selectively dependent on EHMT1/2, whereas H3K9me1 and H3K9me3 are primarily generated independently of these enzymes.

To identify genomic regions marked by H3K9me2, we performed spike-in-normalized chromatin immunoprecipitation followed by sequencing (ChIP-seq) on E14.5 WT and DKO cortices. We identified 15,722 regions decorated by H3K9me2 in the WT cortex, whereas H3K9me2 signal was abolished at the corresponding genomic loci in the DKO cortex (Fig. 1D). H3K9me2 showed minimal overlap with H3K9me3 (Fig. 1E), supporting the conclusion that EHMT1/2 are dispensable for H3K9me3 deposition in the developing cortex.

To characterize the genomic targets of EHMT2, we performed ChIP-seq and identified 11,124 reproducible EHMT2 binding sites in the E14.5 cortex. Of these, 1,023 were associated with promoters, while 9,499 mapped to distal *cis*-regulatory regions within intronic and intergenic sequences (Fig. S2A). EHMT2-bound genomic regions were associated with H3K9me2 but not H3K9me3 in the E14.5 cortex (Fig. 1E). These findings further support that EHMT1/2-mediated H3K9me2 is not a precursor to H3K9me3 and that the two histone marks occupy largely distinct chromatin regions. Furthermore, H3K9me2 showed no overlap with the repressive mark, histone H3 lysine 27 trimethylation (H3K27me3) (Fig. S2B), suggesting that EHMT1/2 operate independently of Polycomb Repressor Complex 2 (PRC2) in the embryonic cortex.

### EHMT1/2 regulate neural progenitor proliferation and the timing of cortical neurogenesis

Loss of *Ehmt1*/2 resulted in a significant reduction in cortical thickness at E15.5, E17.5, and P7 (Fig. 2A). Analysis of cleaved caspase-3 (CC3) during embryonic and postnatal development revealed no increase in apoptotic cells in the DKO cortex (Fig. S3A). To determine whether cortical thinning was associated with defects in neurogenesis, we quantified NSCs and IPCs by immunostaining for PAX6 and TBR2 (EOMES), respectively. The number of PAX6^+^ cells was significantly reduced in E15.5 and E17.5 DKO cortices (Fig. 2B, S3B). Similarly, the number of TBR2^+^ cells was reduced at both stages, suggesting impaired IPC production in the absence of EHMT1/2 (Fig. 2C, S3C). Moreover, the number of cells positive for the proliferation marker Ki67 (MKI67) was significantly reduced in the E15.5 DKO cortex (Fig. 2D). To determine whether this reduction in proliferation was associated with premature cell-cycle exit, we pulse-labeled proliferating cells with 5-ethynyl-2′-deoxyuridine (EdU) at E14.5 and assessed their proliferative status approximately 24 h later (E15.5) by Ki67 staining. The fraction of EdU-labeled cells that had exited the cell cycle (EdU^+^Ki67^-^ cells) was significantly increased in the DKO cortex (Fig. 2E). Together, these results suggest that EHMT1/2 promote the maintenance of a proliferative state and thereby regulate the timing of neural progenitor differentiation.

**Figure 2.**
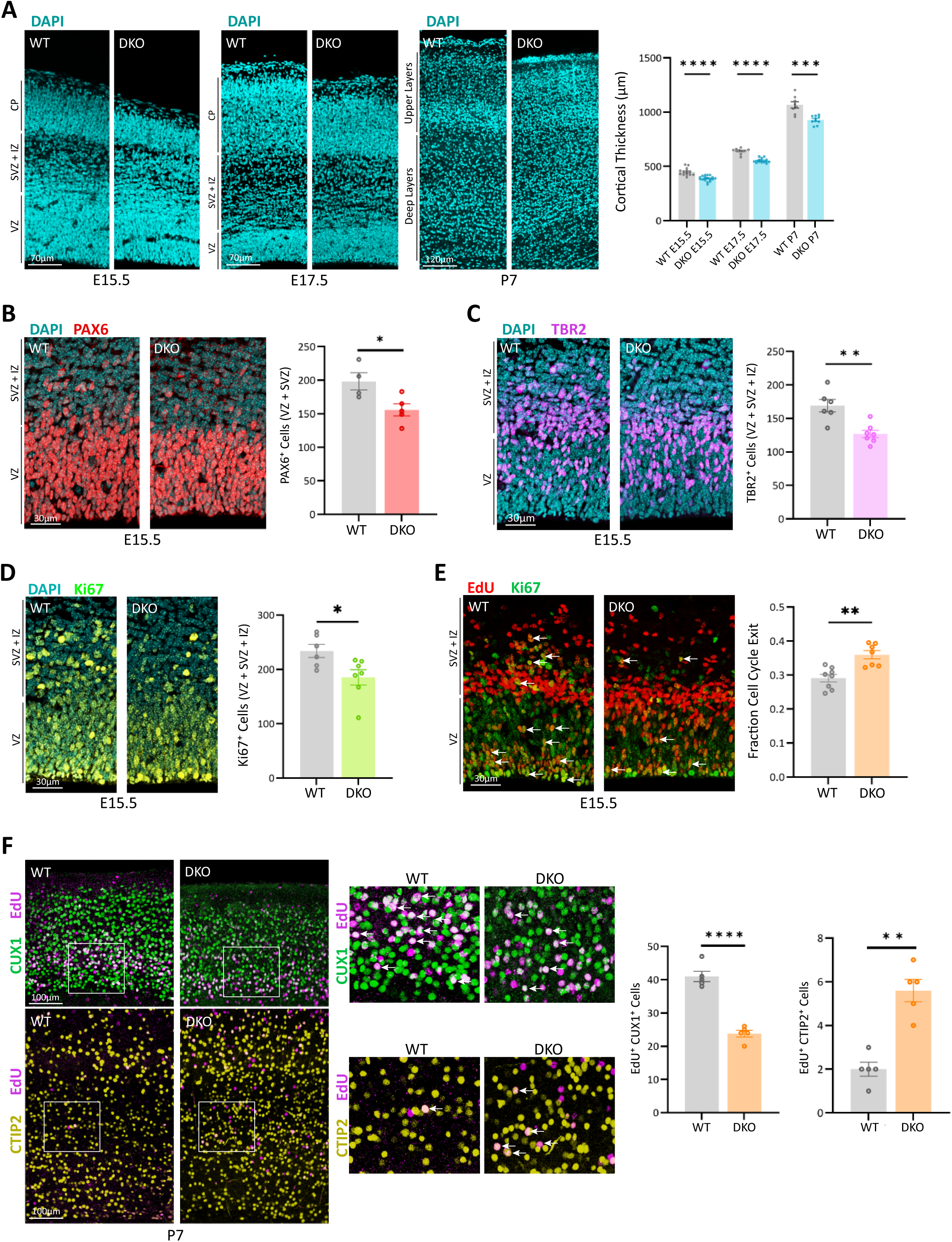
EHMT1 and EHMT2 regulate neurogenesis and the timing of deep-to-upper layer neuron production. **(A)** Coronal sections of the E15.5, E17.5, and P7 cortex showing reduced cortical thickness in the DKO brain (E15.5, n = 14 WT and n = 18 DKO; E17.5, n = 10 WT and n = 15 DKO; P7, n = 8 WT and n = 10 DKO). **(B)** Reduced numbers of PAX6^+^ cells in the E15.5 DKO cortex (n = 4 WT and n = 5 DKO). **(C)** Decreased numbers of TBR2^+^ cells in the E15.5 DKO cortex (n = 6 WT and n = 7 DKO). **(D)** Reduced numbers of Ki67^+^ cells in the E15.5 DKO cortex (n = 6 WT and n = 7 DKO). **(E)** Cell cycle exit analysis following EdU administration at E14.5 and subsequent EdU detection and Ki67 immunostaining approximately 24 h later (E15.5). The DKO cortex exhibited a higher proportion of EdU⁺Ki67⁻ cells, indicating increased cell-cycle exit (n = 8 WT and n = 7 DKO). Accordingly, fewer EdU⁺Ki67^+^cells were observed in the DKO cortex (arrows indicate examples). **(F)** Cell fate analysis following EdU administration at E14.5 and assessment of laminar identity at P7 using the upper-layer marker CUX1 and the deep-layer marker CTIP2. Insets indicate the regions shown at higher magnification. Arrows indicate representative double-labeled cells. The DKO cortex exhibited fewer EdU⁺CUX1⁺ cells and more EdU⁺CTIP2⁺ cells than the WT cortex (n = 5). Quantification was performed within 120-μm-wide cortical columns (B–D) or 150-μm-wide cortical columns (E). VZ, ventricular zone; SVZ, subventricular zone; IZ, intermediate zone; CP, cortical plate. Data are presented as mean ± SEM. Statistical significance was determined using a two-tailed, unpaired Student’s *t*-test; ns, not significant.

We next examined whether loss of *Ehmt1/2* affected cortical layer formation. Analysis of the upper-layer neuronal marker CUX1 and the deep-layer marker CTIP2 in P7 cortices revealed that the overall laminar organization was preserved in DKO mice (Fig. S3D). However, DKO cortices displayed a reduction in upper-layer thickness, significantly fewer CUX1^+^ upper-layer neurons, and an increase in the number of CTIP2^+^ deep-layer neurons (Fig. S3D). These alterations were accompanied by a reduction in the total number of cells within a cortical column, without affecting cell density (Fig. S3D).

To determine whether the altered proportions of upper– and deep-layer neurons in the DKO cortex reflected changes in the temporal progression of neuronal fate specification, we labeled proliferating cells with EdU at E14.5 and subsequently assessed the laminar identities of their neuronal progeny in the P10 cortex. The proportion of EdU-labeled neurons adopting a CTIP2^+^ deep-layer identity was increased in DKO cortices, whereas the proportion acquiring a CUX1^+^ upper-layer identity was decreased (Fig. 2F). Together, these findings indicate that EHMT1/2 regulate neuronal subtype composition in the postnatal cortex by controlling the transition from deep– to upper-layer neurogenesis.

### EHMT1/2 control neuronal differentiation via transcriptional repression and activation

To identify genes dysregulated by the loss of *Ehmt1/2*, we performed single-cell RNA sequencing (scRNA-seq) on E14.5 WT and DKO cortices. This analysis identified 14 transcriptional clusters in both genotypes (Fig. S4A). Based on established marker genes, we assigned 11 clusters to three major cell types along the cortical lineage: NSCs, IPCs, and neurons (Fig. 3A, S4B) (21). We then identified differentially expressed genes (DEGs) in each of these cell types (Fig. 3B, Table S1). A subset of DEGs was shared between neural progenitors and neurons in the DKO cortex, consistent with the broad expression of *Ehmt1*/*2* across the cortical lineage (Fig. 3C).

**Figure 3.**
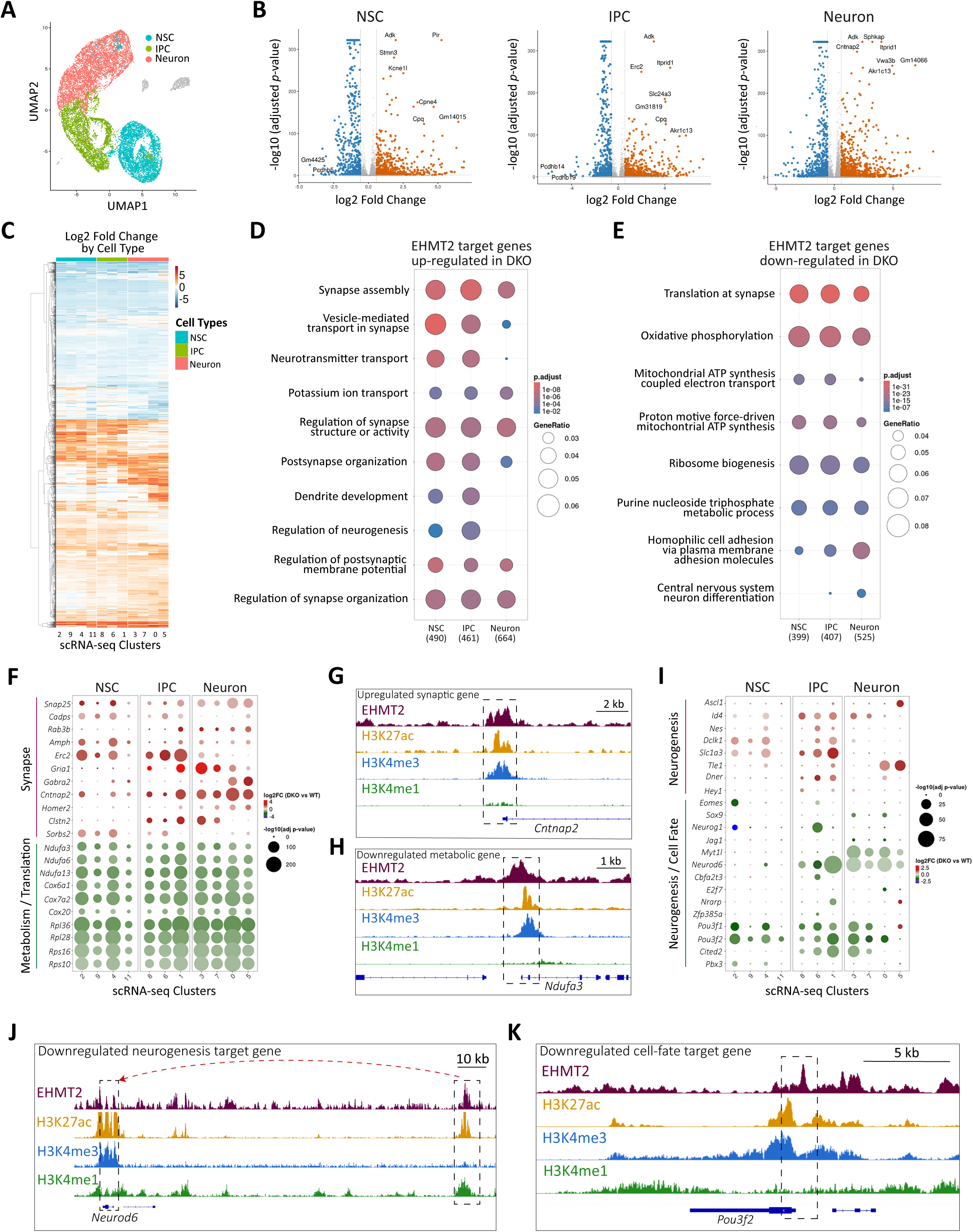
EHMT1 and EHMT2 regulate transcriptional programs of neurogenesis and cortical lineage progression. **(A)** UMAP plot showing the three major cell subtypes identified by scRNA-seq in the E14.5 cortex: Neural stem cell (NSC), intermediate progenitor cell (IPC), and neuron. **(B)** Volcano plots showing differentially expressed genes between WT and DKO NSCs, IPCs, and neurons. **(C)** Heatmap showing relative fold changes in gene expression values between WT and DKO cortices along distinct stages of neurogenesis. Numbers below the heatmap indicate the corresponding scRNA-seq clusters for each cell type. **(D, E)** Gene Ontology (GO) analysis of predicted EHMT2 target genes that were upregulated (D), consistent with repression by EHMT2, or downregulated (E), consistent with activation by EHMT2, in the DKO cortex across NSCs, IPCs, and neurons. Numbers in parentheses below the dot plots indicate the total number of genes in each cell type that mapped to significantly enriched GO terms. **(F)** Dot plot showing fold changes and *p*-values for selected synaptic and metabolic target genes across individual NSCs, IPCs, and neuron scRNA-seq clusters. **(G, H)** Representative genomic tracks showing EHMT2 binding at the promoter of the synaptic gene *Cntnap2* (G) and the metabolic gene *Ndufa3* (H). **(I)** Dot plot showing fold changes and *p*-values for selected genes associated with neurogenesis and upper-layer neuron specification across individual NSCs, IPCs, and neuron scRNA-seq clusters. **(J, K)** Representative genomic tracks showing EHMT2 binding at an enhancer of the neurogenesis-associated gene *Neurod6* (J) and EHMT2 binding at the promoter of the upper-layer specification gene *Pou3f2* (K). H3K27ac, H3K4me3, and H3K4me1 ChIP-seq tracks from the E14.5 cortex are included in G, H, J, and K.

The transcriptional changes observed in the DKO cortex could reflect either direct regulation by EHMT1/2 or indirect downstream effects due to their loss. To distinguish between these possibilities, we integrated the scRNA-seq data with EHMT2 ChIP-seq profiles (Fig. S5A-C). DEGs in the DKO cortex whose promoters were bound by EHMT2 were classified as direct proximal targets (Fig. S5A-C). To identify potential distal targets of EHMT2, we further incorporated previously published high-throughput chromosome conformation capture (3C) with sequencing (Hi-C) datasets characterizing enhancer–promoter interactions in the embryonic mouse cortex (Fig. S5A-C) (22). This integrative analysis identified 521 EHMT2-bound promoters in the WT cortex, 249 of which were associated with genes that were either upregulated or downregulated in the DKO cortex (Table S2). In addition, among 1419 EHMT2-bound enhancers with predicted target genes assigned based on Hi-C interactions, 442 were linked to promoters of either upregulated or downregulated genes (Table S2). Together, these analyses identified candidate proximal and distal target genes of EHMT2 and revealed its association with both transcriptional repression and activation in the embryonic cortex.

### EHMT1/2 primarily regulate genes associated with synapse formation and metabolism during corticogenesis

Gene Ontology (GO) analysis of upregulated target genes in the DKO cortex revealed significant enrichment for synapse-associated genes (Fig. 3D). Notably, loss of *Ehmt1/2* resulted in the premature activation of synaptic genes in NSCs and IPCs (Fig. 3D, Table S2). EHMT2 directly targeted genes involved in synaptic vesicle exocytosis, including *Snap25*, *Cadps*, and *Erc2*; synaptic scaffolding, including *Homer2* and *Clstn2*; and synaptic adhesion, including *Cntnap2* (Fig. 3F) (23). For example, EHMT2 occupied the *Cntnap2* promoter in the WT cortex (Fig. 3G), whereas *Cntnap2* expression was increased in the DKO cortex (Fig. 3F). These findings suggest that EHMT1/2 restrain synaptic gene expression in neural progenitors and neurons, thereby regulating the timing of synaptic gene activation during cortical neurogenesis and neuronal maturation.

In contrast, downregulated target genes in the DKO cortex were primarily associated with metabolic processes (Fig. 3E). Loss of *Ehmt1/2* reduced the expression of *Ndufa* and *Cox* genes, which encode subunits of mitochondrial respiratory chain Complex I (NADH dehydrogenase) and Complex IV (cytochrome c oxidase), respectively, and contribute to oxidative phosphorylation (Fig. 3E, F) (24). Downregulated targets were also enriched for genes associated with translation, a category composed predominantly of ribosomal genes from the *Rpl* and *Rps* families (Fig. 3E, F, Table S2) (25). Notably, EHMT2 occupied the promoters of several metabolic and ribosomal genes identified as potential direct targets (Fig. 3H, S6A, Table S2). These findings suggest that EHMT2 activates genes involved in oxidative phosphorylation and ribosome biogenesis during cortical neurogenesis.

### EHMT1/2 regulate neurogenesis and cell fate specification genes in the embryonic cortex

We next sought to identify EHMT2 target genes that might contribute to the microcephaly and reduced production of upper-layer neurons observed in DKO cortices. Among the DEGs, we identified a subset associated with neurogenesis and cell fate specification (Fig. 3I). Several neurogenesis-associated genes, including *Ascl1*, *Id4*, *Nes*, *Dclk1*, *Slc1a3*, and *Hey1*, were upregulated in the DKO cortex, and EHMT2 occupied the promoters and/or enhancers of these genes in the WT cortex, consistent with direct transcriptional repression (Fig. 3I, S6B, Table S2). Conversely, another subset of neurogenesis-associated target genes, including *Tbr2*, *Sox9*, *Neurog1*, *Jag1*, *Mytl1*, and *Neurod6*, was downregulated in the DKO cortex, consistent with direct activation by EHMT2 in the WT cortex (Fig. 3I, J, S6B, Table S2).

We also identified reduced expression of genes encoding key determinants of upper-layer neuronal identity, including *Pou3f1* (*Oct6*), *Pou3f2* (*Brn2*), *Pou3f3* (*Brn1*), *Cited2*, and *Pbx3*, in the DKO cortex (Fig. 3I) (26–28). EHMT2 occupancy at the regulatory regions of these genes in the WT cortex was consistent with direct transcriptional activation (Fig. 3I, K, S6B). Importantly, the expression of upper-layer neuron identity genes was reduced in neural progenitors (Fig. 3I), potentially biasing their neuronal progeny toward a deep-layer identity and contributing to the reduced production of upper-layer neurons in DKO cortices. Collectively, these findings suggest that EHMT1/2 regulate cortical growth and laminar composition through the coordinated repression and activation of genes involved in neurogenesis and neuronal fate specification.

### EHMT2 controls gene expression through H3K9me2-dependent and –independent mechanisms

To distinguish EHMT2 functions dependent on H3K9me2 from those independent of this mark, we classified the ChIP-seq datasets into three distinct genomic-coordinate clusters. Cluster 1 (625 sites) exhibited high EHMT2 binding and high H3K9me2 enrichment (Fig. 4A, D). Cluster 2 (10,499 sites) displayed high EHMT2 binding with low to undetectable H3K9me2 levels (Fig. 4B, E). Cluster 3 (15,101 sites) showed high H3K9me2 enrichment and low EHMT2 binding (Fig. 4C, F).

**Figure 4.**
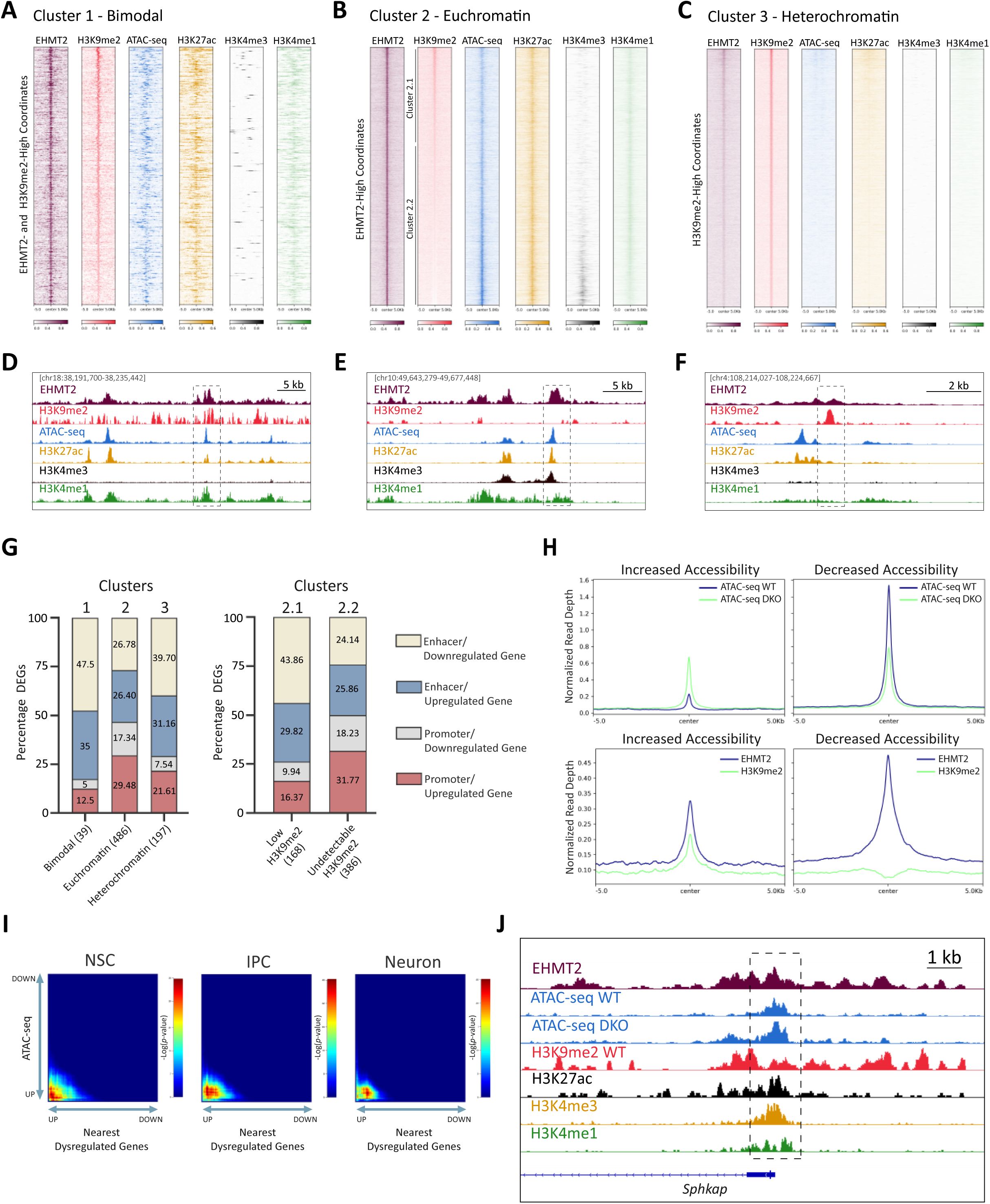
EHMT2 regulates neurogenesis through distinct chromatin modalities. **(A)** Cluster 1 comprises genomic coordinates with high EHMT2 binding and high H3K9me2 enrichment. This cluster is predominantly composed of enhancers that display a “bimodal” chromatin signature, characterized by a repressive mark (H3K9me2) overlapping regions with high accessibility and active enhancer marks (H3K27ac and H3K4me1). **(B)** Cluster 2 comprises genomic regions with high EHMT2 binding. These regions exhibit features of euchromatin, including high accessibility and active histone marks. Cluster 2 was further subdivided into cluster 2.1, showing low H3K9me2, and cluster 2.2, showing minimal to undetectable H3K9me2. **(C)** Cluster 3 comprises genomic regions with high H3K9me2 enrichment. This cluster displays low EHMT2 binding. The absence of chromatin accessibility and active histone marks indicates that cluster 3 predominantly represents heterochromatin. Within each heatmap, genomic regions are ordered according to mean H3K9me2 signal intensity, with the highest signal at the top. For all heatmaps, the DKO H3K9me2 signal was subtracted from the WT signal to reduce background and improve signal specificity. **(D-F)** Representative genomic tracks illustrating the distinct EHMT2 and H3K9me2 signatures corresponding to clusters 1-3, respectively (dashed boxes). **(G)** Stacked bar plots showing the percentage of differentially expressed genes (DEGs) in the DKO cortex associated with *cis*-regulatory elements assigned to clusters 1-3. Color-coded sections distinguish promoter and enhancer elements associated with either upregulated or downregulated genes. Numbers in parentheses along the *x*-axis indicate the total number of DEGs associated with each cluster. A total of 59 DEGs showed both promoter and enhancer interactions and were therefore included in two categories. **(H)** Projection profiles showing ATAC-seq signal in WT and DKO cortices (*top*), together with EHMT2 occupancy and H3K9me2 enrichment (*bottom*), across genomic regions exhibiting increased (*left*) or decreased (*right*) chromatin accessibility. **(I)** Rank-rank hypergeometric overlap analysis showing the relationship between differential chromatin accessibility and differential gene expression in NSCs, IPCs, and neurons from the E14.5 DKO cortex. Low *p*-values in the bottom-left corner of each heatmap indicate a significant correlation between increased chromatin accessibility and upregulated expression of the nearest genes. Arrows indicate the positions of genomic regions with increased or decreased accessibility (*y*-axis) and upregulated or downregulated genes (*x*-axis). **(J)** Representative genomic tracks showing EHMT2 binding at the promoter of the synaptic gene *Sphkap* (dashed box). ATAC-seq tracks show increased chromatin accessibility at this region in the DKO compared with the WT cortex. EHMT2 binding overlaps with H3K9me2.

Cluster 1 exhibited high chromatin accessibility, as determined by the Assay for Transposase-Accessible Chromatin using sequencing (ATAC-seq), together with elevated levels of the active histone marks H3K27ac and H3K4me1 (Fig. 4A, D), consistent with these regions representing enhancers (29). We classified this cluster as bimodal, as H3K9me2 overlaps with active histone marks, possibly reflecting heterogeneity in the chromatin states of these enhancers (Fig. 4A, D). Cluster 2 comprised regions with high chromatin accessibility and enrichment for active enhancer (H3K27ac and H3K4me1) and active promoter marks (H3K27ac and H3K4me3) (Fig. 4B, E). Lastly, cluster 3 comprised regions of inaccessible chromatin lacking active histone marks (Fig. 4C, F). Together, these profiles reveal distinct patterns of EHMT2 binding and H3K9me2 levels in the embryonic cortex. Cluster 1 consists of heterogeneous chromatin states, cluster 2 primarily encompasses euchromatin, and cluster 3 corresponds to heterochromatin.

To link these distinct EHMT2 and H3K9me2 profiles to gene regulation, we intersected the genomic coordinates of the three clusters with *cis*-regulatory elements linked to DEGs in the DKO cortex (Fig. 4G). More DEGs were associated with promoters and/or enhancers mapping to cluster 2 (486) compared with cluster 3 (197), suggesting that most genes regulated by EHMT2 are within the euchromatin compartment (Fig. 4G). We further subdivided cluster 2 into two subclusters based on H3K9me2 levels: cluster 2.1 comprised regions with low H3K9me2, whereas cluster 2.2 encompassed regions with minimal to undetectable H3K9me2 (Fig. 4B). Of the 486 DEGs associated with cluster 2, 168 had regulatory elements linked to cluster 2.1, while 386 had regulatory elements linked to cluster 2.2 (Fig. 4G), suggesting that EHMT2 regulates most genes within euchromatin largely independently of H3K9me2 in the embryonic cortex.

To investigate how EHMT1/2 loss affects chromatin accessibility, we performed ATAC-seq on E14.5 WT and DKO cortices. We identified 1,047 regions with increased accessibility and 2,646 regions with decreased accessibility in the DKO cortex (Fig. 4H). Most regions that gained accessibility in the DKO cortex were bound by EHMT2 and marked by H3K9me2 in the WT cortex (Fig. 4H), consistent with a role for H3K9me2 in promoting chromatin compaction (30). Conversely, most regions that lost accessibility in the DKO cortex were bound by EHMT2 but lacked detectable H3K9me2 in the WT cortex (Fig. 4H), suggesting that these regions are associated with EHMT2-mediated transcriptional activation. Consistent with their distinct accessibility profiles, regions that gained accessibility in the DKO cortex exhibited low levels of active histone marks in the WT cortex, indicating that these regions are maintained in a predominantly repressed state by EHMT2 (Fig. S7A). In contrast, regions that lost accessibility in the DKO cortex were enriched for active histone marks in the WT cortex (Fig. S7A). Collectively, these findings suggest that EHMT2 regulates gene expression through distinct mechanisms: H3K9me2-associated chromatin compaction at repressed regulatory elements and H3K9me2-independent maintenance of high accessibility at transcriptionally active regulatory elements.

We next investigated whether changes in chromatin accessibility were associated with altered gene expression in the DKO cortex by comparing differential ATAC-seq peaks with the nearest DEGs using rank-rank hypergeometric overlap (RRHO) analysis (31). Regions that gained accessibility were strongly associated with upregulated genes in NSCs, IPCs, and neurons in the DKO cortex (Fig. 4I). In contrast, the association between regions that lost accessibility and downregulated genes did not reach statistical significance at any stage of differentiation (Fig. 4I). Moreover, upregulated genes associated with increased chromatin accessibility were enriched for synaptic functions (Fig. 4J, S7B). Notably, a distinct group of synaptic genes repressed by EHMT1/2 was not associated with increased chromatin accessibility in the DKO cortex and mapped to cluster 2.2, which exhibited minimal to undetectable H3K9me2 levels (Fig. 4B, S7B). Collectively, these findings suggest that EHMT2 restrains the premature activation of synaptic genes in neural progenitors and neurons through both H3K9me2-dependent chromatin compaction and H3K9me2-independent regulation.

### EHMT1/2 cooperate with PRDM16 to repress synaptic genes independently of H3K9me2

To further elucidate the mechanisms by which EHMT1/2 regulate gene expression, we sought to identify their transcriptional partners in the embryonic cortex. Previous studies have demonstrated that EHMT1/2 interact with the chromatin regulator PRDM16 (positive regulatory domain-containing protein 16) in adipocytes to inhibit muscle differentiation (32, 33). In the E14.5 cortex, *Ehmt1/2* and *Prdm16* expression overlapped in NSCs and a subset of IPCs (Fig. 5A). Co-immunoprecipitation of PRDM16 in cultured cortical neural progenitors confirmed its interaction with both EHMT1 and EHMT2 (Fig. 5B). Consistent with this interaction, a subset of EHMT2 and PRDM16 binding sites overlapped in the embryonic cortex (Fig. 5C-E). Most co-occupied regions showed low to undetectable H3K9me2 (Fig. 5C, D), suggesting that EHMT1/2 and PRDM16 cooperate in gene regulation largely independently of H3K9-methylated heterochromatin.

**Figure 5.**
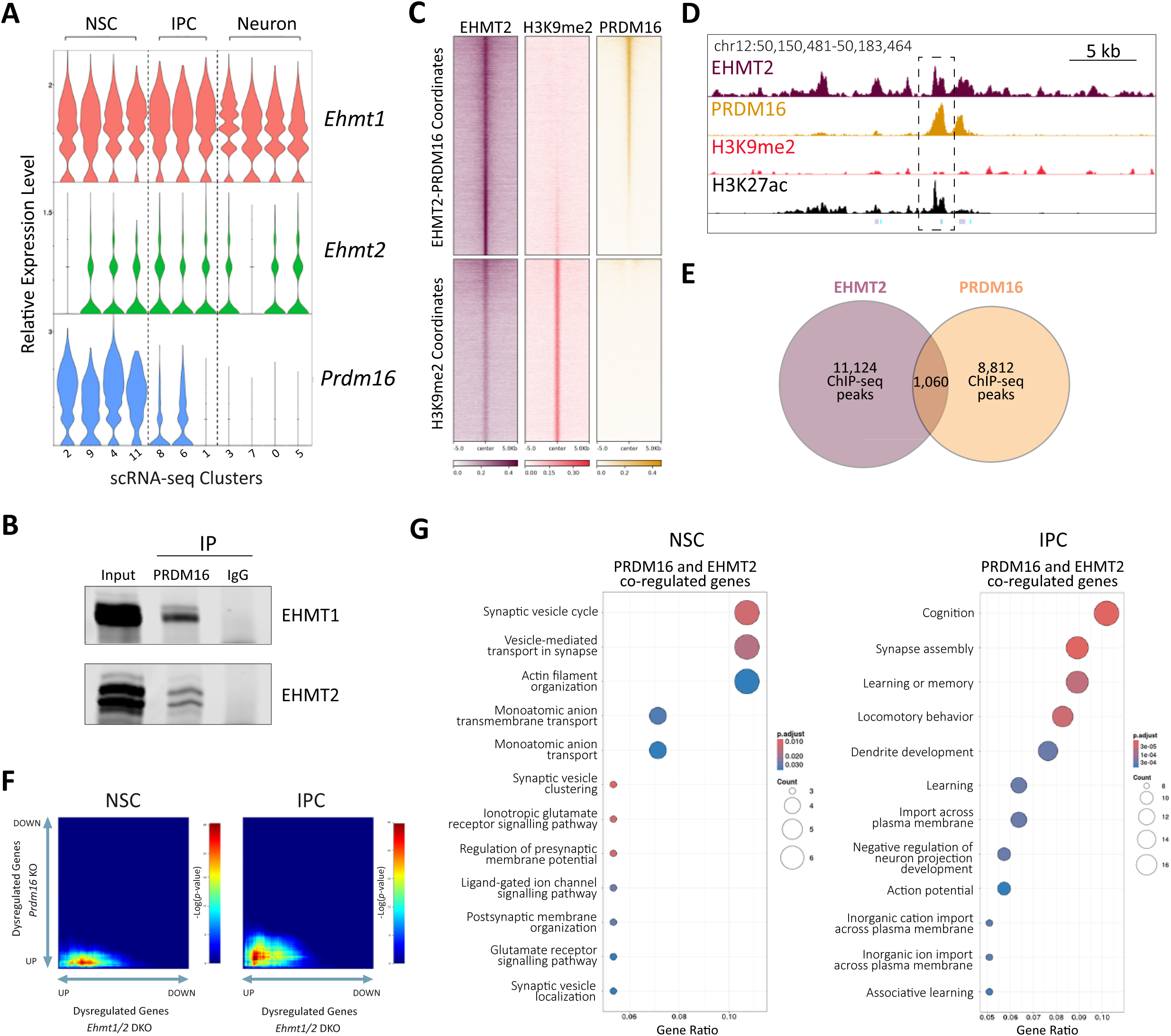
EHMT1 and EHMT2 cooperate with PRDM16 to repress synaptic genes in neural progenitors. **(A)** Violin plots showing single-cell expression levels of *Ehmt1*, *Ehmt2*, and *Prdm16* in the E14.5 cortex. Numbers below the plots indicate the corresponding scRNA-seq clusters for each cell type. **(B)** Co-immunoprecipitation of PRDM16 followed by western blotting demonstrates its association with EHMT1 and EHMT2 in cultured neural progenitors isolated from the E14.5 cortex. Nonspecific IgG was used as a negative control. The blot is representative of two independent co-immunoprecipitation experiments. **(C)** Heatmaps showing EHMT2, H3K9me2, and PRDM16 ChIP-seq signal across the union of EHMT2 and PRDM16 peak coordinates (top panel) and H3K9me2 coordinates (bottom panel). Genomic regions are ordered according to mean PRDM16 signal intensity, with the highest signal at the top. **(D)** Representative genomic tracks showing colocalization of EHMT2 and PRDM16 at an enhancer element exhibiting H3K27ac enrichment but lacking detectable H3K9me2 (dashed box). **(E)** Venn diagrams showing the overlap between EHMT2 and PRDM16 ChIP-seq peaks in the embryonic cortex. **(F)** Rank-rank hypergeometric overlap analysis comparing dysregulated genes between *Ehmt1/2* DKO and *Prdm16* KO NSCs and IPCs. Arrows indicate the positions of upregulated and downregulated genes within each heatmap. Low *p*-values in the bottom-left corner of each heatmap indicate a significant correlation between genes upregulated in both knockout models, consistent with shared gene repression by EHMT1/2 and PRDM16. **(G)** Gene Ontology analysis of biological processes associated with genes co-repressed by EHMT1/2 and PRDM16 in NSCs and IPCs indicates enrichment for synaptic functions.

To test whether EHMT1/2 and PRDM16 co-regulate gene expression during cortical neurogenesis, we compared DEGs in NSCs and IPCs from *Ehmt1/2* DKO and *Prdm16* SKO cortices (34). A subset of upregulated and downregulated genes overlapped between the two genotypes in NSCs and IPCs (Fig. S7C). We next performed RRHO analysis to identify genes co-regulated by EHMT1/2 and PRDM16. This analysis identified a prominent group of genes that were upregulated following the loss of either EHMT1/2 or PRDM16 in both NSCs and IPCs, consistent with a cooperative role of these regulators in gene repression (Fig. 5F). These co-repressed genes were strongly enriched for synapse-associated functions in both NSCs and IPCs (Fig. 5G). Together, these findings suggest that EHMT1/2 cooperate with PRDM16 to repress synaptic genes independently of H3K9me2, thereby preventing their premature activation in neural progenitors during cortical development.

## DISCUSSION

Here, we demonstrate that EHMT1/2 play a specific role in establishing H3K9me2 in the embryonic cortex, with minimal impact on H3K9me1 levels and no apparent effect on H3K9me3. This finding contrasts with *Ehmt1/2*-null mouse embryos or cell lines, in which H3K9me1 levels were substantially reduced (10, 11). These differences indicate that H3K9me1 may be established predominantly by non-canonical H3K9 methyltransferases in the developing cortex (35). Our data further indicate that EHMT1/2 account for most H3K9me2 in the embryonic cortex, with no evidence of substantial EHMT1/2-independent H3K9me2 deposition, as described in embryonic stem cells *in vitro* (9).

A previous study reported that EHMT1/2 associate with the H3K9 methyltransferases SETDB1 and SUV39H1 in cell lines (36). These interactions have supported a model in which H3K9 methyltransferases cooperate in transcriptional silencing through the sequential generation of H3K9me2 by EHMT1/2, followed by H3K9me3 deposition by SETDB1, SUV39H1, and SUV39H2 (36). In contrast, our findings indicate that H3K9me2 and H3K9me3 are established through distinct mechanisms and occupy separate genomic regions in the embryonic cortex. Consistent with this interpretation, we recently showed that combined loss of SETDB1, SUV39H1, and SUV39H2 selectively depletes H3K9me3 in the embryonic cortex and induces transcriptional and chromatin alterations distinct from those observed following EHMT1/2 depletion and H3K9me2 loss (37). Despite these differences, depletion of either H3K9me2 or H3K9me3 in the developing cortex results in microcephaly, although the phenotype is more severe following H3K9me3 depletion (37). Thus, H3K9me2 and H3K9me3 appear to define separate chromatin landscapes and regulate cortical neurogenesis—and, ultimately, cortical size—through mechanistically distinct pathways.

Our results indicate that EHMT1/2 regulate the balance between neural progenitor proliferation and differentiation, as well as the timing of cortical neuron specification, by directly modulating the expression of genes involved in these processes through transcriptional repression and activation. Moreover, loss of *Ehmt1/2* in the developing cortex results in premature activation of synaptic genes in neural progenitors, which may further contribute to the accelerated neurogenesis observed in the DKO cortex (38). Consistent with this possibility, recent evidence has implicated EHMT1/2 in restraining neuronal differentiation and maturation in embryonic stem cell-derived human neurons *in vitro* (38). In addition, EHMT1/2 promote the expression of genes associated with oxidative phosphorylation and ribosome biogenesis in the developing cortex. In this context, neural progenitors primarily rely on aerobic glycolysis to support proliferation, whereas differentiating neurons increasingly depend on mitochondrial oxidative phosphorylation to meet their elevated energy demands (39). Collectively, these findings indicate that EHMT1/2 play a central role in coordinating neurogenic and synaptic transcriptional programs with the metabolic and biosynthetic pathways that support neuronal differentiation and synaptic function.

Our analysis revealed distinct patterns of EHMT2 occupancy and H3K9me2 enrichment that define different chromatin environments in the embryonic cortex. Among these, we identified a subset of enhancers characterized by enrichment of the active marks H3K4me1 and H3K27ac alongside the repressive mark H3K9me2 (cluster 1). Typically, enhancer-associated bivalent chromatin is characterized by the absence of H3K27ac and the coexistence of H3K4me1 with the repressive mark H3K27me3 and, less frequently, H3K9me3 (29, 40). Thus, cluster 1 enhancers are unlikely to represent bivalent or poised elements but may instead correspond to dynamically regulated enhancers whose activity varies across cell types. We also identified genomic regions with high EHMT2 occupancy but low to undetectable H3K9me2 (cluster 2), which were predominantly associated with euchromatin, and regions with low EHMT2 occupancy but high H3K9me2 (cluster 3), which were primarily associated with heterochromatin. These distinct chromatin profiles suggest that clusters 1 and 2 participate in dynamic gene regulation, whereas cluster 3 may comprise genomic regions associated with more stable transcriptional repression. Furthermore, the constitutive heterochromatin profile of cluster 3 raises the possibility that these regions are enriched at the nuclear periphery and overlap with LADs, where they may reside in a transcriptionally repressive environment (6). Future studies will be needed to determine whether loss of EHMT1/2 and H3K9me2 alters the spatial organization of these genomic regions within nuclear compartments during cortical neurogenesis.

Our data suggest that EHMT2 regulates distinct subsets of target genes through H3K9me2-dependent and –independent mechanisms. For example, a subset of EHMT2-repressed synaptic genes was associated with H3K9me2 at their promoters and/or enhancers. These genes exhibited increased accessibility following EHMT1/2 loss in the DKO cortex, supporting a role for H3K9me2 in chromatin compaction and transcriptional repression (30). In contrast, a distinct subset of synaptic genes appeared to be repressed independently of H3K9me2, as EHMT2 occupied their regulatory regions in the absence of detectable H3K9me2. Thus, H3K9me2-independent activities of EHMT2, such as promoting DNA methylation, could contribute to this repression (41). Similarly, most metabolic genes whose expression was positively associated with EHMT2 occupancy lacked detectable H3K9me2. Together, these findings suggest that EHMT1/2 regulate subsets of synaptic and metabolic genes independently of H3K9me2, potentially enabling rapid and coordinated changes in gene expression during neuronal differentiation.

This study implicates PRDM16 as a transcriptional partner through which EHMT1/2 regulate gene expression independently of H3K9me2 in cortical neural progenitors. This finding was unexpected, given that PRDM16 has been reported to catalyze H3K9me1 *in vitro* and to contribute to the formation of H3K9me2-marked heterochromatin through its interaction with EHMT1/2 in cultured adipocytes (32, 33, 42). In contrast, we did not detect significant H3K9me2 enrichment at PRDM16-bound regions in the embryonic cortex, suggesting that the EHMT1/2–PRDM16 complex may operate through a distinct mechanism in this context. PRDM16 has also been implicated in regulating the migration of upper-layer neurons through interactions with CtBP1/2 and the recruitment of HDAC1/2 to enhancers (34, 43). Although EHMT1/2 have also been reported to interact with CtBP1/2 (44), we did not observe neuronal migration defects in the DKO cortex. This difference may reflect the partial overlap between EHMT2– and PRDM16-bound *cis*-regulatory regions in the developing cortex, leading to both shared and distinct downstream transcriptional targets. Moreover, our analysis identified a subset of synaptic genes as prominent candidate targets of coordinated repression by EHMT1/2 and PRDM16. These findings support a model in which the EHMT1/2–PRDM16 complex represses synaptic genes in neural progenitors, thereby preventing their premature expression before neuronal differentiation. As neural progenitors differentiate into neurons, downregulation of *Prdm16* may relieve this repression, facilitating the activation of synaptic gene expression in newly generated neurons.

Loss of EHMT1 in the embryonic mouse cortex resulted in a reduction of EHMT2 and H3K9me2 in neural progenitors comparable to that observed in the DKO cortex. This finding indicates that EHMT1 loss compromises EHMT2 expression or stability, thereby exacerbating the depletion of H3K9me2. These results raise the possibility that *EHMT1* haploinsufficiency in humans similarly reduces EHMT2 abundance or activity, possibly contributing to the molecular and developmental abnormalities associated with Kleefstra syndrome (16, 18). In addition, the DKO microcephaly parallels the reported microcephaly in Kleefstra patients (17), suggesting that *EHMT1* haploinsufficiency might also lead to deficits in neurogenesis and altered neuronal subtype composition in humans.

## MATERIAL AND METHODS

### Mouse strains

*Ehmt1^lox/lox^* mice carry *lox*P sites flanking exon 23, which encodes part of the histone methyltransferase SET domain (13). *Ehmt2^lox/lox^* mice harbor *lox*P sites flanking exons 23 and 24, which also encode a portion of the SET domain (19). Cre-mediated recombination results in nonsense-mediated decay of the mutant *Ehmt2* transcript (19). *Ehmt1^lox/lox^*, *Ehmt2^lox/lox^* mice were mated with *Emx1^Cre^* mice (B6.129S2-Emx1tm1(cre)Krj/J, strain #005628-JAX) to produce cortex-specific DKO animals (20). All mice used to generate primary cell cultures were ICR (CD-1) from Envigo. All housing, husbandry, EdU treatment, and euthanasia procedures were performed in accordance with protocols approved by the Bloomington Institutional Animal Care and Use Committee (BIACUC) at Indiana University. Mouse sex was not determined before experimentation; however, mouse embryo sex was retrospectively inferred for the scRNA-seq analysis.

### Immunostaining and EdU labeling

Embryonic and postnatal brains were fixed in 4% paraformaldehyde (PFA), sectioned using a VT1000 vibratome (Leica Biosystems), and incubated in antigen retrieval solution (10 mM sodium citrate, 0.05% Tween 20, pH 6.0) for 1 h at 70°C. Sections were incubated in blocking buffer (10% goat serum, 0.1% Triton X-100, 0.01% sodium azide in 1X PBS) followed by overnight incubation with primary antibodies diluted in fresh blocking buffer. After washing in 1X PBS, sections were incubated in secondary antibodies diluted in blocking buffer for 2 h. Nuclei were counterstained with DAPI (4′,6-diamidino-2-phenylindole). The following primary antibodies were used: anti-H3K9me1 (Abcam, ab8896, 1:1000), anti-H3K9me2 (Abcam, ab1220, 1:1000), anti-H3K9me3 (Active Motif, 39062, 1:1000), anti-EHMT1 (Abcam, ab41969, 1:500), anti-EHMT2 (Abclonal A19288, 1:500), anti-Ki67 (Abcam, ab16667, 1:250), anti-cleaved caspase-3 (Cell Signaling Technology, 9661, 1:2000), anti-PAX6 (BioLegend, 901301, 1:300), anti-TBR2 (Invitrogen, 14-4875-82, 1:250), anti-CUX1 (Millipore, ABE217, 1:500), and anti-CTIP2 (Abcam, ab18465, 1:500). We used the following secondary antibodies from Invitrogen: Alexa Fluor 488 anti-mouse (A11001) and anti-rabbit (A11008); Alexa Fluor 546 anti-mouse (A11003), anti-rabbit (A11010), and anti-rat (A11081); Alexa Fluor 647 anti-rabbit (A21244) and anti-rat (A21247), all diluted 1:500. EdU^+^ cells were detected using the Click-iT EdU Imaging Kit with Alexa Fluor 647 (Invitrogen, C10340). Fluorescent images were acquired using a Leica SP8 confocal microscope (Leica Microsystems).

### Primary cortical NSC cultures

E14.5 WT cortices were pooled and dissociated into a single-cell suspension by incubation in 0.05% trypsin–EDTA (Gibco, 25300062) containing 0.01% DNase (Sigma-Aldrich, DN-25). Following enzymatic digestion, the trypsin solution was removed, and cells were washed twice with calcium-free Hibernate-E medium (BrainBits, HECA500). Tissue was then mechanically dissociated in the same medium using a fire-polished Pasteur pipette. Cells were resuspended in complete NSC medium composed of Dulbecco’s Modified Eagle Medium/Nutrient Mixture F12 (DMEM/F12; Gibco, 11320033), 1X GlutaMax (Gibco, 35050-061), 1X B27, 10 ng/ml murine Epidermal Growth Factor (EGF; PeproTech, 315-09), 10 ng/ml human Fibroblast Growth Factor 2 (FGF2; PeproTech, 100-18), and 0.12 mg/ml bovine serum albumin (Gibco, 15260037). Cells were plated onto culture dishes pre-coated with 10 μg/ml Poly-D-Lysine (Corning, 354210) and 12 μg/ml Laminin (Gibco, 23017015).

### Immunoprecipitation

To extract nuclear proteins, frozen pellets of neural progenitors expanded *in vitro* were resuspended in hypotonic Buffer A (10 mM HEPES pH 7.5, 1 mM MgCl₂, 10 mM KCl, 0.5 mM DTT) containing 1X protease inhibitor (Thermo Scientific, 87786). Cells were mechanically disrupted on ice using a pestle, followed by centrifugation to pellet nuclei. The nuclear pellet was resuspended in extraction Buffer B (20% glycerol, 20 mM HEPES, pH 7.3, 250 mM NaCl, 1.5 mM MgCl₂, 0.2 mM EDTA, 0.5 mM DTT, and 1X protease inhibitor) and further homogenized with a pestle until the suspension became cloudy, confirming nuclear lysis and chromatin release. The lysate was centrifuged at 20,000 *g* for 10 min at 4°C, and the supernatant containing the nuclear extract was recovered. Immunoprecipitation was conducted by incubating nuclear extracts overnight at 4°C with antibody-conjugated Protein G Dynabeads (Invitrogen, 10003D). After incubation, beads were sequentially washed with Buffer B, high salt wash buffer (20 mM HEPES pH 7.3, 500 mM NaCl, 1.5 mM MgCl2, 0.2 mM EDTA, 10% glycerol, 0.1% NP40), and low salt wash buffer (20 mM HEPES pH 7.3, 150 mM NaCl, 1.5 mM MgCl2, 0.2 mM EDTA, 10% glycerol, 0.1% NP40). Immunoprecipitated proteins were eluted with elution buffer (0.1 M glycine, 0.1% NP40 pH 2.5, and 1X protease inhibitor) and immediately neutralized with 1 M Tris, pH 8. To enhance immunoprecipitation efficiency, cortical neural progenitors *in vitro* were infected with a retrovirus expressing full-length *Prdm16* (43). A sheep anti-PRDM16 antibody (R&D Systems, AF6295) was used for immunoprecipitation.

### Western blot

Protein samples were resolved on 4–20% mini-Protean TGX precast gels (Bio-Rad), transferred to PVDF membranes, and incubated in blocking buffer (3% Bovine Serum Albumin, 20 mM Tris-HCl pH 7.5, 150 mM NaCl, and 0.1% Tween 20) for 1 h. Membranes were incubated with primary antibodies overnight at 4°C, washed three times with TBST 1X (Tris-buffered saline with Tween 20), and incubated with secondary antibodies for 1 h, followed by three additional washes with TBST 1X. The following antibodies were used: rabbit anti-EHMT1 (Abclonal, A8513, 1:1000), rabbit anti-EHMT2 (Cell Signaling Technology. 3306, 1:1000), sheep anti-IgG control antibody (Millipore, 12-370), anti-rabbit IgG secondary antibody (LICORbio, 926-32211, 1:5000 dilution), anti-sheep IgG HRP-conjugated secondary antibody (R&D Systems, HAF016, 1:1000).

### Single-cell RNA-seq

E14.5 WT and DKO cortices were pooled and dissociated into single-cell suspensions as described above for primary cultures. Cells were resuspended at approximately 1,200 cells/μl in calcium– and magnesium-free DPBS (Gibco, 14190144) supplemented with 0.04% BSA (Sigma, A3059). Cell viability was assessed by Trypan Blue exclusion, and visual inspection confirmed the absence of cell doublets. Single-cell RNA-seq (scRNA-seq) libraries were prepared using the Chromium Next GEM Single Cell 3′ Reagent Kits v3.1 (10x Genomics, PN-100269) and the Dual Index Kit TT Set A (10x Genomics, PN-1000215). Libraries were sequenced on a NovaSeq X Series instrument using the 1.5B Reagent Kit (Illumina, 20104703).

### EHMT2 ChIP-seq

Cortices from an entire E14.5 WT CD-1 litter were pooled and homogenized using a pestle, followed by sequential dual crosslinking with 1.5 mM ethylene glycol bis(succinimidyl succinate) (EGS; Thermo Scientific) for 20 min and subsequently with 1% paraformaldehyde (PFA) for 10 min. Crosslinking was quenched with 125 mM glycine, and samples were washed twice with 1X PBS plus protease inhibitor (Thermo Scientific, 87786). Cells were lysed in lysis buffer (20 mM Tris-HCl pH 8.0, 85 mM KCl, 0.5% NP40) and nuclei were resuspended in SDS buffer (0.2% SDS, 20 mM Tris-HCl pH 8.0, 1 mM EDTA). Chromatin was sheared using a Covaris S220 ultrasonicator to generate predominantly mono– and dinucleosomal fragments. Fragmented chromatin was diluted 1:1 with 2X ChIP dilution buffer (0.1% sodium deoxycholate, 2% Triton X-100, 2 mM EDTA, 30 mM Tris-HCl pH 8.0, 300 mM NaCl) and incubated overnight at 4°C with 10 µl (21 µg/ml) of anti-EHMT2 antibody (C6H3 Cell Signaling Technology). Washed Protein G Dynabeads (Invitrogen, 10003D) were added to the samples and incubated for 2 h at 4°C. Beads were collected and sequentially washed with low salt wash buffer (0.1% SDS, 1%Triton X-100, 2 mM EDTA, 20 mM Tris-HCl pH 8.0, 150 mM NaCl), high salt wash buffer (0.1% SDS, 1% Triton X-100, 2 mM EDTA, 20 mM Tris-HCl pH 8.0, 500 mM NaCl), LiCl wash buffer (0.25 M LiCl, 0.5% NP40, 0.5% sodium deoxycholate, 1 mM EDTA, 10 mM Tris-HCl pH 8.0), and TE buffer (10 mM Tris-HCl pH 8.0, 1 mM EDTA). Chromatin was eluted using elution buffer (1% SDS, 0.1 M NaHCO3) and reverse-crosslinked overnight at 65°C in reverse-crosslinking buffer (250 mM Tris-HCl, pH 6.5; 62.5 mM EDTA, pH 8.0; 1.25 M NaCl; 5 mg/ml proteinase K). DNA was extracted with phenol/chloroform/isoamyl alcohol and precipitated with 3 M sodium acetate, pH 5.0. Samples were resuspended in TE buffer (10 mM Tris-HCl, pH 8.0, 0.1 mM EDTA) supplemented with RNase A (100 mg/ml) and incubated for 1 h at 37°C. Samples were purified using the DNA Clean & Concentrator Magbead Kit (Zymo Research, D4012), and sequencing libraries were prepared using the NEBNext Ultra II DNA Library Prep kit (NEB, E7103S) with NEBNext Multiplex Oligos for Illumina (NEB, E7335S). Libraries were loaded on a P2 (100 Cycles) flow cell configured to produce 2×61 nucleotides paired-end reads and sequenced on a NextSeq 2000 instrument.

### H3K9me2 ChIP-seq

Individual E14.5 cortices were homogenized with a pestle in 100 μl nuclear extraction buffer (10 mM Tris-HCl pH 8.0, 140 mM NaCl, 5 mM MgCl_2_, 0.6% NP40), yielding approximately 2.5 to 3.5 million nuclei per sample. Isolated nuclei were digested with micrococcal nuclease (MNase) using a master mix prepared per ChIP sample by combining 15 μl 10X MNase buffer (NEB, M0247S), 2.2 μl dithiothreitol (DTT; 100 mM stock, Thermo Scientific, 707265ML), 1.5 μl halt protease inhibitor (100X stock, Thermo Scientific, 87785), 1.5 μl phenylmethylsulfonyl fluoride (PMSF; 100 mM stock, Thermo Scientific, 36978), 10 μl MNase (2 × 10^6^ U/ml stock, NEB, M0247S), and nuclease-free water to a final volume of 50 μl. MNase digestion was performed by adding 50 μl of the master mix to each sample and incubating for 12 min at 37°C. Reactions were stopped by the addition of 1/10 volume of 100 mM EDTA, followed by 1/10 volume of 1% Triton X-100 and 1% sodium deoxycholate. Chromatin was diluted with complete immunoprecipitation buffer (20 mM Tris-HCl pH 8.0, 2 mM EDTA, 150 mM NaCl, 0.1% Triton X-100, 1X halt protease inhibitor, 1X PMSF) supplemented with 18 ng spike-in chromatin (Active Motif, 53083). Samples were precleared by incubation with 50 µl pre-washed Protein G Dynabeads (Invitrogen, 1003D) for 2-6 h at 4°C with rotation. Precleared chromatin was incubated overnight at 4°C with antibody-Protein G bead complexes containing 5 μg anti-H3K9me2 antibody (Abcam, ab1220) and 2 μg of spike-in antibody (Active Motif, 61686) per ChIP sample. Beads were washed sequentially with complete immunoprecipitation buffer, low salt buffer (20 mM Tris-HCl pH 8.0, 2 mM EDTA, 150 mM NaCl, 1% Triton X-100, 0.1% SDS, 1X halt protease inhibitor), and high salt buffer (20 mM Tris-HCl pH 8.0, 2 mM EDTA, 500 mM NaCl, 1% Triton X-100, 0.1% SDS, 1X halt protease inhibitor). Chromatin was eluted using freshly prepared elution buffer (0.1 M NaHCO_3,_ 1% SDS), and RNA was removed by adding 4 μl 1 M Tris-HCl (pH 6.5), 4 μl 0.5 M EDTA, and 2 μl RNase A (stock at 20 μg/μl). DNA was extracted with phenol/chloroform/isoamyl alcohol and precipitated with 3 M sodium acetate, pH 5.0. DNA pellets were washed with 70% ethanol and resuspended in TE buffer (10 mM Tris-HCl, pH 8.0, 0.1 mM EDTA). Samples were further purified using the DNA Clean & Concentrator Magbead Kit (Zymo Research, D4012). ChIP-seq libraries were prepared using the Ultra II DNA library Prep Kit for Illumina (NEB, E7103S) and NEBNext Multiplex Oligos for Illumina (NEB, E7335S). Spike-in–normalized H3K9me2 ChIP-seq from the *Ehmt1/2* DKO cortex was used as a negative control.

### ATAC-seq

Aliquots of approximately 50,000 cells were prepared from E14.5 WT and DKO cortices as described above for primary cultures. ATAC-seq libraries were prepared using the Zymo-Seq ATAC Library Kit (Zymo, D5458) according to the manufacturer’s instructions. Library size distribution was determined using a TapeStation 2200 system (Agilent) before sequencing, as described above for ChIP-seq.

### Bioinformatic analyses

#### RNA-seq analysis

FASTQ files from scRNA-seq were processed using Cell Ranger v9.0.1 from 10x Genomics (https://www.10xgenomics.com/). Embryonic sex was inferred based on expression of Y-chromosome genes across WT and DKO barcodes. Barcodes showing expression of *Uty*, *Eif2s3y*, *Ddx3y*, and *Kdm5d* were identified as male. To eliminate sex-specific transcriptional bias, all male barcodes from WT and DKO datasets were excluded. Downstream analyses were conducted with Seurat v5.3.1 (45) using R v4.5.1. After QC filtering, data were normalized with SCTransform, integrated across samples, and subjected to principal component analysis. Cell clusters were visualized using uniform manifold approximation and projection (UMAP). Marker genes for each cluster were identified with the FindMarkers function, enabling annotation of NSCs, IPCs, and cortical neurons. Differential gene expression was assessed using the Wilcoxon test implemented in FindMarkers, applying thresholds of *q*-value ≤ 0.05 and abs(log2FC) ≥ 0.58. Gene ontology analysis was conducted using the R packages clusterProfiler 4.18.4 (46) and enrichplot 1.30.5 (DOI: 10.18129/B9.bioc.enrichplot.)

Previously reported bulk RNA-seq gene expression counts in isolated NSCs and IPCs from *Prdm16* WT and KO mouse lines were included (34). DESeq2 v1.48.2 was used to perform differential gene expression analysis for each cell type (47).

#### H3K9me2 ChIP-seq analysis

The H3K9me2 ChIP-seq experiment was normalized for immunoprecipitation efficiency using Drosophila melanogaster spike-in chromatin (48). The normalized raw reads were trimmed using fastp v1.0.1 (49) and aligned to a combined mouse and Drosophila reference genome using bowtie2 v2.5.4 (50). Mouse– and Drosophila-derived reads were separated using split_bam.py from the Spiker package v1.0.5 (51). Duplicate reads were removed using MarkDuplicates from the Picard Toolkit v3.4.0, and unpaired reads were excluded with reformat.sh from BBTools v39.52 (https://bbmap.org). Read counts for each sample were scaled by a factor equal to the minimum number of dm6-mapped reads across all samples divided by the number of dm6-mapped reads for that sample (48). Peak calling was conducted with MACS3 v3.0.3 (52) for WT samples using combined DKO samples as negative controls with “-c” parameter and a *q*-value cutoff of 0.05. Blacklisted genomic regions were removed using BEDTools v2.31.1 (53, 54). Peak reproducibility was determined by requiring 30% overlap between replicates to report a peak.

#### EHMT2 and PRDM16 ChIP-seq analysis

EHMT2 and the previously reported PRDM16 ChIP-seq raw reads were trimmed using fastp v1.0.1 and aligned to the mm39 reference genome using HISAT2 v2.2.1 (34, 49, 55). Reads mapping to mitochondrial DNA (ChrM), ‘ChrX’, ‘ChrY’, or with a MAPQ score <2 were removed using SAMtools v1.23 (56). Duplicate reads were removed using MarkDuplicates from the Picard Toolkit v3.4.0 (https://broadinstitute.github.io/picard/; Broad Institute). bigWig files were generated using bamCoverage from deepTools v3.5.6 with cpm normalization (57). Peak calling was performed using MACS3 v3.0.3 with “-q 0.05” except for EHMT2 replicate 1, which used a relaxed cutoff of “-p 1e-2” (52). Blacklisted genomic regions were removed from the BED files using BEDTools v2.31.1 (53, 54). Peak reproducibility was determined by requiring at least 30% overlap between replicates. EHMT2 reproducible peaks were assigned to genomic features using annotatePeaks.pl from the HOMER suite v5.1 with the default Homer mm39 annotation (58). This method of associating EHMT2 peaks to promoters was only used to characterize the population of peaks, and it is distinct from the method described below to identify EHMT2 target genes.

#### Identification of EHMT2 target genes

To identify genes directly regulated by EHMT2, we intersected the list of DEGs in the DKO cortex with the EHMT2 ChIP-seq data. The genes displaying EHMT2 binding at their annotated promoters, which were defined as ±1.5 Kb from the transcription start site of dysregulated genes, were identified as proximal EHMT2 targets. To identify distal EHMT2 target genes, we merged the Hi-C coordinates of enhancer-promoter pairs in neural progenitor and cortical neurons and converted them from mm10 to mm39 using the UCSC LiftOver tool (22). We then identified EHMT2-bound enhancers that overlapped these enhancer–promoter pairs and linked them to DEGs in the DKO cortex. The associated genes were classified as distal EHMT2 targets.

#### Integration of EHMT2 and H3K9me2 ChIP-seq

EHMT2 and H3K9me2 peaks were partitioned into three clusters using the bedtools intersect command. Cluster 1 comprised EHMT2 peaks that overlapped H3K9me2 peaks by at least 30%. Cluster 2 comprised the remaining EHMT2 peak regions, and Cluster 3 comprised the remaining H3K9me2 peak regions. Cluster 2 exhibited strong EHMT2 binding across all regions, while a subset of these showed weak H3K9me2 signal that did not meet the statistical threshold for peak calling. However, this weak H3K9me2 signal was reduced in the DKO samples, suggesting that it reflects genuine H3K9me2 enrichment. Cluster 2 was further subdivided into subclusters 2.1 and 2.2 by applying k-means clustering to binned EHMT2 intensity values generated using the computeMatrix and plotHeatmap tools from the deepTools package (57). Cluster 3 exhibited strong H3K9me2 enrichment accompanied by a weak EHMT2 signal. To determine whether the weak EHMT2 signal in Cluster 3 represented specific EHMT2 binding, we generated 10,000 randomized permutations of the Cluster 3 peak set using the bedtools shuffle command while preserving the number and lengths of the peaks. We then used multiBigwigSummary to calculate the mean EHMT2 signal intensity across the genomic regions in each permutation. The observed mean EHMT2 signal intensity in Cluster 3 was compared with the distribution of mean signal intensities from the randomized sets. An empirical *p*-value was calculated as *p* = (*r* + 1)/(*n* + 1), where *r* is the number of randomized sets with a mean EHMT2 signal intensity equal to or greater than that observed in Cluster 3, and *n* is the total number of permutations. The resulting *p*-value was 0.0001, indicating that the weak EHMT2 signal in Cluster 3 was significantly higher than expected by chance, consistent with specific EHMT2 binding within these regions.

#### Other ChIP-seq datasets

Previously reported H3K4me1, H3K4me3, H3K27ac, H3K9me3, and H3K27me3 ChIP-seq datasets were included (37, 59, 60). Mapped reads from H3K27me3 ChIP-seq in NSC and neuron samples were merged using SAMtools, and a CPM-normalized BigWig file was generated using bamCoverage. The UCSC LiftOver tool was used to convert peaks called against mm9 to mm39 and the merge tools in BEDTools was used to merge NSC and neuron peaks.

#### ATAC-seq analysis

Raw reads were trimmed using fastp v1.0.1 and aligned to the mm39 reference genome using HISAT2 v2.2.1 (49, 55). Reads mapping to ‘ChrM’, ‘ChrX’, ‘ChrY’, or with a MAPQ score <2 were removed using SAMtools v1.23 (56). Duplicate reads were removed using MarkDuplicates from the Picard Toolkit v3.4.0 (https://broadinstitute.github.io/picard/; Broad Institute). bigWig files were generated using bamCoverage from deepTools v3.5.6 with cpm normalization (57). Peak calling was performed using MACS3 v3.0.3 with relaxed cutoff “-p 1e-2” (52). Blacklisted genomic regions were removed from the BED files using the BEDTools v2.31.1 (53, 54). Peak reproducibility was assessed using IDR v2.0.4.2 (61). Differential accessibility analysis was performed using DiffBind v3.18.0 (62).

#### Rank-Rank Hypergeometric Overlap analysis

The correlation between differential gene expression and differential chromatin accessibility was computed with the RRHO test as implemented in the R package RRHO 1.48.0 (31). Differentially accessible regions were assigned to the nearest gene using annotatePeaks.pl from the HOMER suite v5.1 (58). RRHO analysis was also performed to compare the DKO scRNA-seq with previously published bulk RNA-seq data from *Prdm16* KO NSCs and IPCs isolated from the E15.5 mouse cortex (34).

### Statistical analysis

#### Image quantification

Histone methylation immunostaining was quantified using single-plane confocal images converted to 16-bit format in ImageJ, with the default pixel intensity threshold applied. Regions of interest (ROIs) were manually defined based on the DAPI channel overlaid with the immunostaining channel, and pixel intensities were measured in 30–50 ROIs per condition. To account for variability in signal intensity across experiments, pixel intensities were normalized using the following formula:

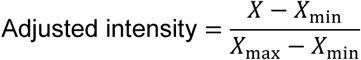

where X represents the histone methylation signal intensity within a nucleus, and X_min_ and X_max_ represent the minimum and maximum intensities, respectively, observed across WT and DKO samples.

#### Cortical thickness, cell number, and cell density quantification

Cortical thickness was measured in anatomically equivalent regions of the somatosensory cortex as the distance from layer I to the white matter tracts using LAS X Life Science software (Leica Microsystems). Cell counts were obtained either manually using ImageJ/Fiji or with the LAS X cell counter software. At least three brains per genotype, derived from a minimum of two litters, were analyzed. Cell density was calculated by dividing the total number of DAPI-stained nuclei by the corresponding cortical area.

#### Cell-cycle exit quantification

The cell-cycle exit index was calculated as the proportion of EdU⁺ Ki67⁻ cells among all EdU⁺ cells within a cortical column of defined width, approximately 24 h after EdU injection.

In all quantifications, at least three brains per genotype from a minimum of two litters were analyzed per condition. Statistical significance was assessed using a two-tailed, unpaired Student’s *t*-test in GraphPad Prism. Significance thresholds were defined as \**p* < 0.05, \*\**p* < 0.01, \*\*\**p* < 0.001, and \*\*\*\**p* < 0.0001.

## Acknowledgements

We are grateful to Dr. Anne Schaefer (Icahn School of Medicine) and Dr. Alexander Tarakhovsky (The Rockefeller University) for the kind gift of the *Ehmt1^lox/lox^* and *Ehmt2^lox/lox^* mouse lines. We thank David Merritt, Jun Yin, and Reza Abdi at the Center for Genomics and Bioinformatics (Indiana University Bloomington) for their valuable support with next-generation sequencing experiments.

## Author contributions

Conceptualization: J.-M.B.; Investigation: A.C., S.K., J.-M.B.; Methodology: A.C., J.-M.B.; Validation: A.C., S.K., J.-M.B.; Software: C.H., R.P.; Data curation: A.C., C.H., R.P.; Formal analysis: A.C., C.H., R.P.; Visualization: A.C., C.H., R.P..; Funding acquisition: J.-M.B.; Project administration: J.-M.B.; Supervision: J.-M.B.; Writing – original draft: J.-M.B.; Writing – review & editing: A.C., C.H., J.-M.B.

## Data availability

The raw sequencing data generated in this study have been deposited in the NCBI Sequence Read Archive (SRA) under accession number PRJNA1531914.

## Supplemental information

**Document S1**. Figures S1–S7 with legends and supplementary table legends.

**Table S1. DEGs between E14.5 WT and DKO cortices.** This table lists DEGs identified by scRNA-seq in NSCs, IPCs, and cortical neurons. Each tab lists the DEGs identified in the corresponding cell type.

**Table S2. Predicted EHMT2 target genes in the E14.5 cortex.** Tab 1 lists predicted proximal targets with EHMT2-bound promoters, and Tab 2 lists predicted distal targets associated with EHMT2-bound enhancers. Each tab indicates the subset of differentially expressed targets in the DKO cortex.

## Supplementary Figure Legends

**Figure S1.**
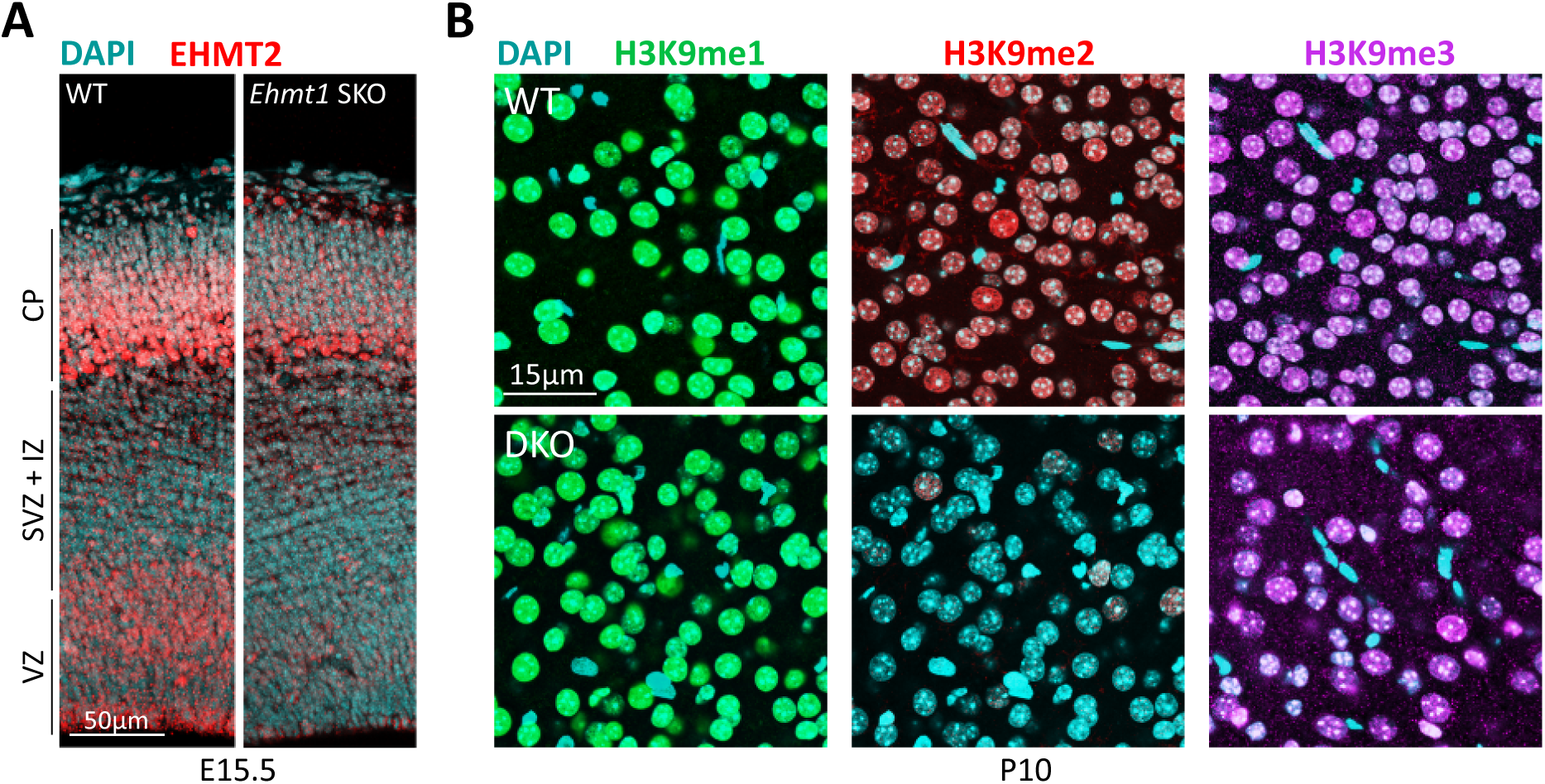
(**A**) Immunostaining showing a pronounced reduction in EHMT2 signal in the VZ and SVZ + IZ of the *Ehmt1* SKO cortex. **(B)** Immunostaining of the P10 cortex showing H3K9me2 depletion in DKO cortical neurons, whereas H3K9me1 and H3K9me3 remained largely unchanged. VZ, ventricular zone; SVZ, subventricular zone; IZ, intermediate zone; CP, cortical plate.

**Figure S2.**
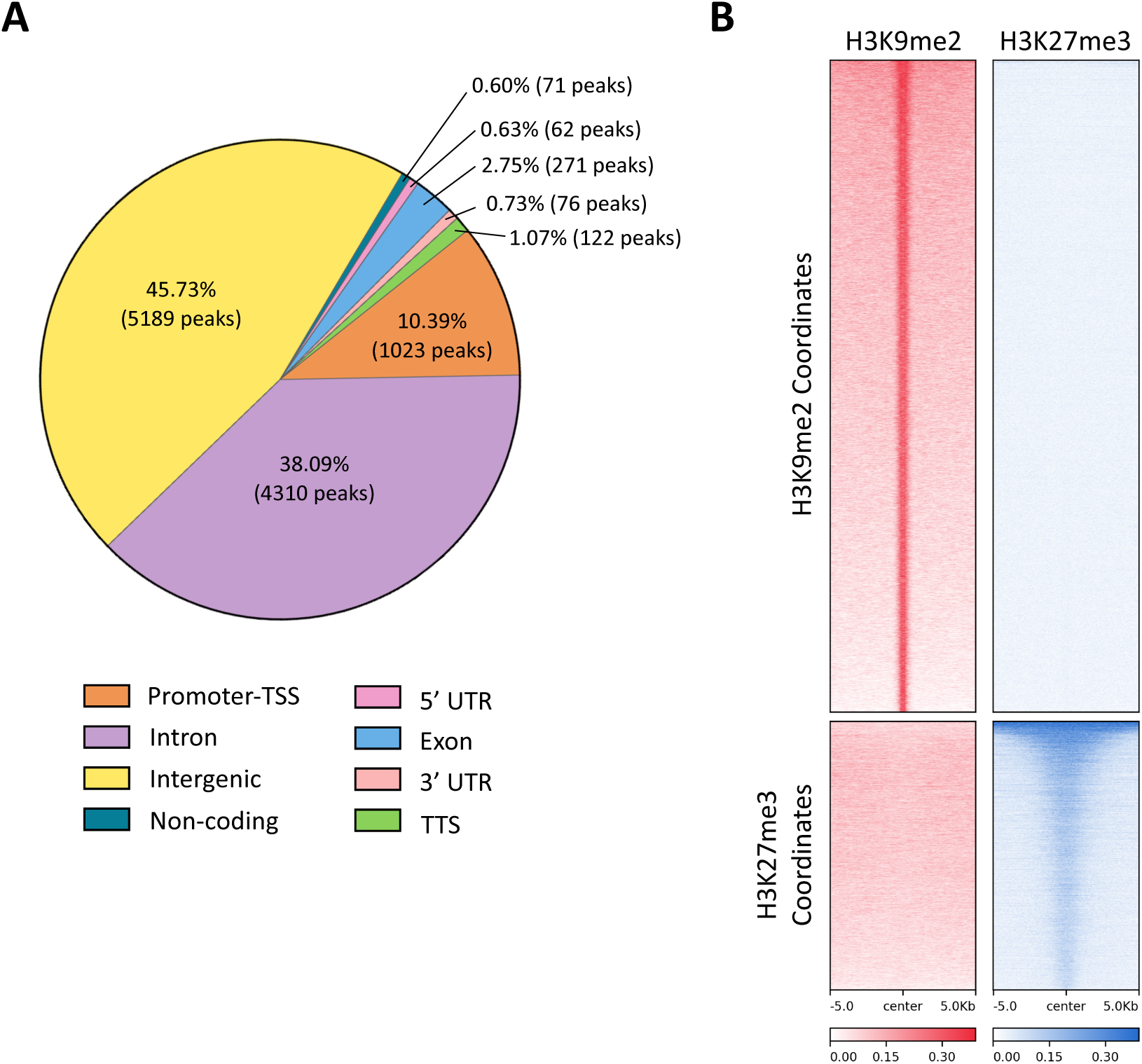
(**A**) Pie chart showing the distribution of EHMT2 ChIP-seq peaks across genomic features in the E14.5 cortex. TSS, transcription start site; TTS, transcription termination site; UTR, untranslated region. **(B)** H3K9me2 and H3K27me3 ChIP-seq heatmaps across genomic coordinates centered on H3K9me2 peaks (top) or H3K27me3 peaks (bottom), showing minimal overlap between these histone modifications in the E14.5 cortex. Heatmaps were sorted by mean signal intensity across samples, with the highest signal at the top.

**Figure S3.**
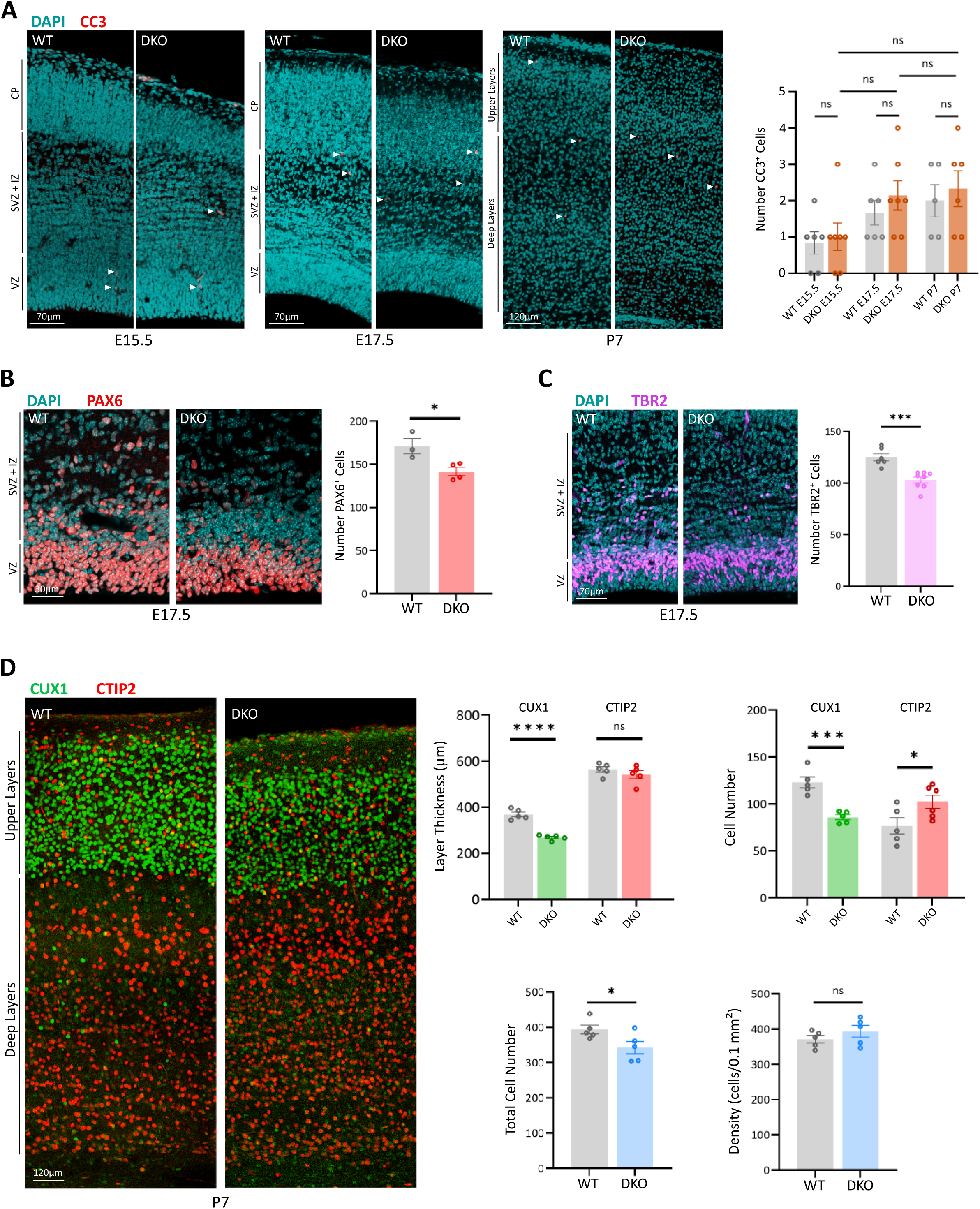
(**A**) Coronal sections of WT and DKO cortices immunostained for cleaved caspase-3 (CC3) at E15.5, E17.5, and P7. Arrowheads indicate representative CC3⁺ cells. No significant differences in CC3⁺ cell numbers were observed between genotypes (E15.5, n = 6 WT and n = 7 DKO; E17.5, n = 6 WT and n = 7 DKO; P7, n = 5 WT and n = 6 DKO). **(B, C)** PAX6 and TBR2 immunostaining showing a significant reduction in PAX6⁺ (panel B; n = 3 WT and n = 4 DKO) and TBR2⁺ (panel C; n = 6 WT and n = 8 DKO) cells in the E17.5 DKO cortex. **(D)** Coronal sections of P7 WT and DKO cortices immunostained for CTIP2, a deep-layer neuronal marker, and CUX1, an upper-layer neuronal marker. The upper plots show quantification of upper– and deep-layer cortical thickness based on CUX1 and CTIP2 staining, respectively (n = 5), as well as the number of CUX1⁺ and CTIP2⁺ neurons (n = 5). The lower plots show quantification of total cell numbers (n = 5) and cell density (n = 5) based on DAPI staining. Quantification was performed within 120-μm-wide cortical columns (B, C) or 100-μm-wide cortical columns (D). VZ, ventricular zone; SVZ, subventricular zone; IZ, intermediate zone; CP, cortical plate. Data are presented as mean ± SEM. Statistical significance was determined using two-tailed, unpaired Student’s *t*-tests; ns, not significant.

**Figure S4.**
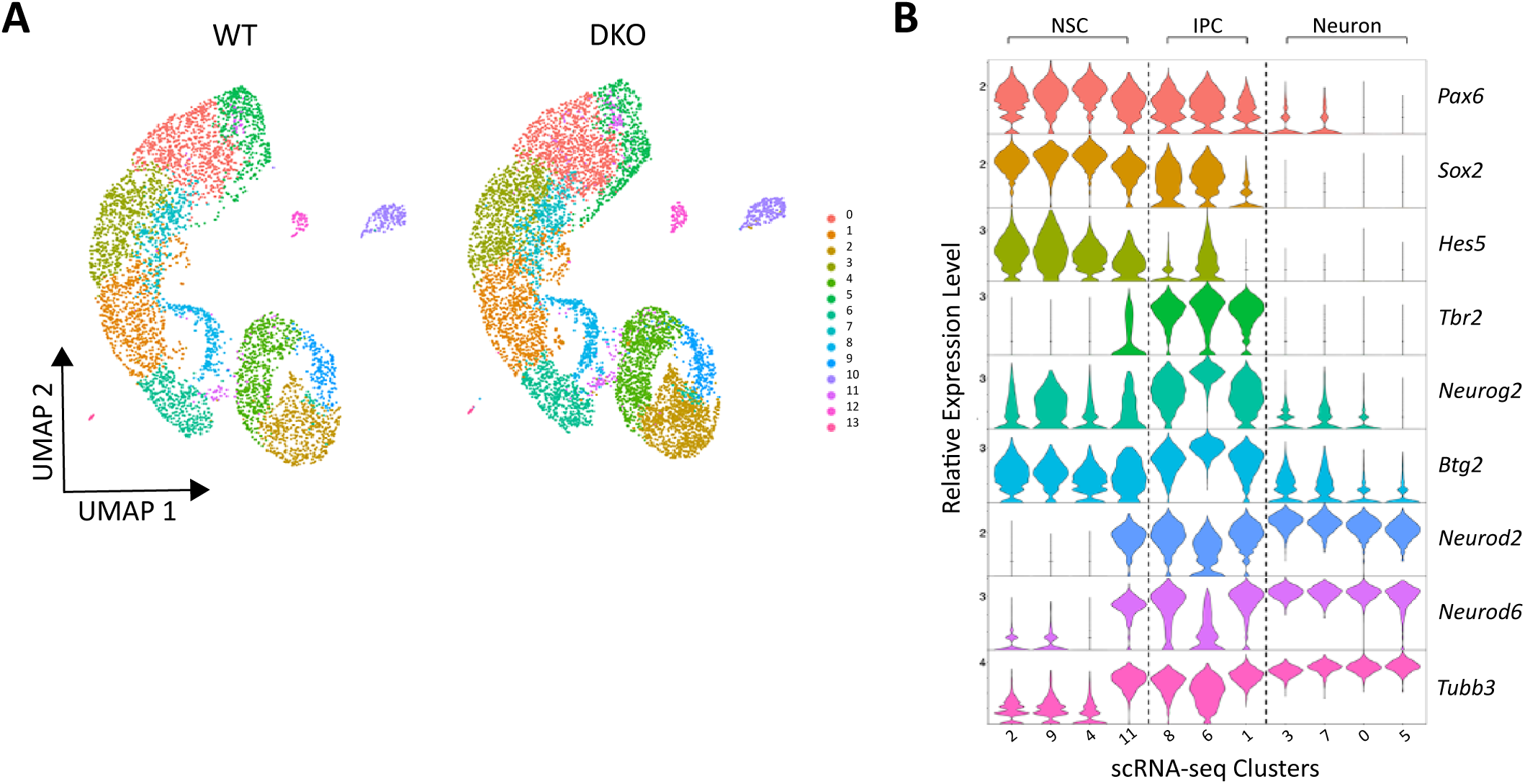
(**A**) UMAP plots showing an equivalent distribution of corresponding scRNA-seq clusters in E14.5 WT and DKO cortices. **(B)** Violin plots showing the expression of established cell-type markers for NSCs (*Pax6*, *Sox2*, and *Hes5*), IPCs (*Tbr2*, *Neurog2*, and *Btg2*), and cortical neurons (*Neurod2*, *Neurod6*, and *Tubb3*). Numbers below the violin plots indicate the 11 transcriptional clusters shown in panel A corresponding to NSCs, IPCs, and neurons.

**Figure S5.**
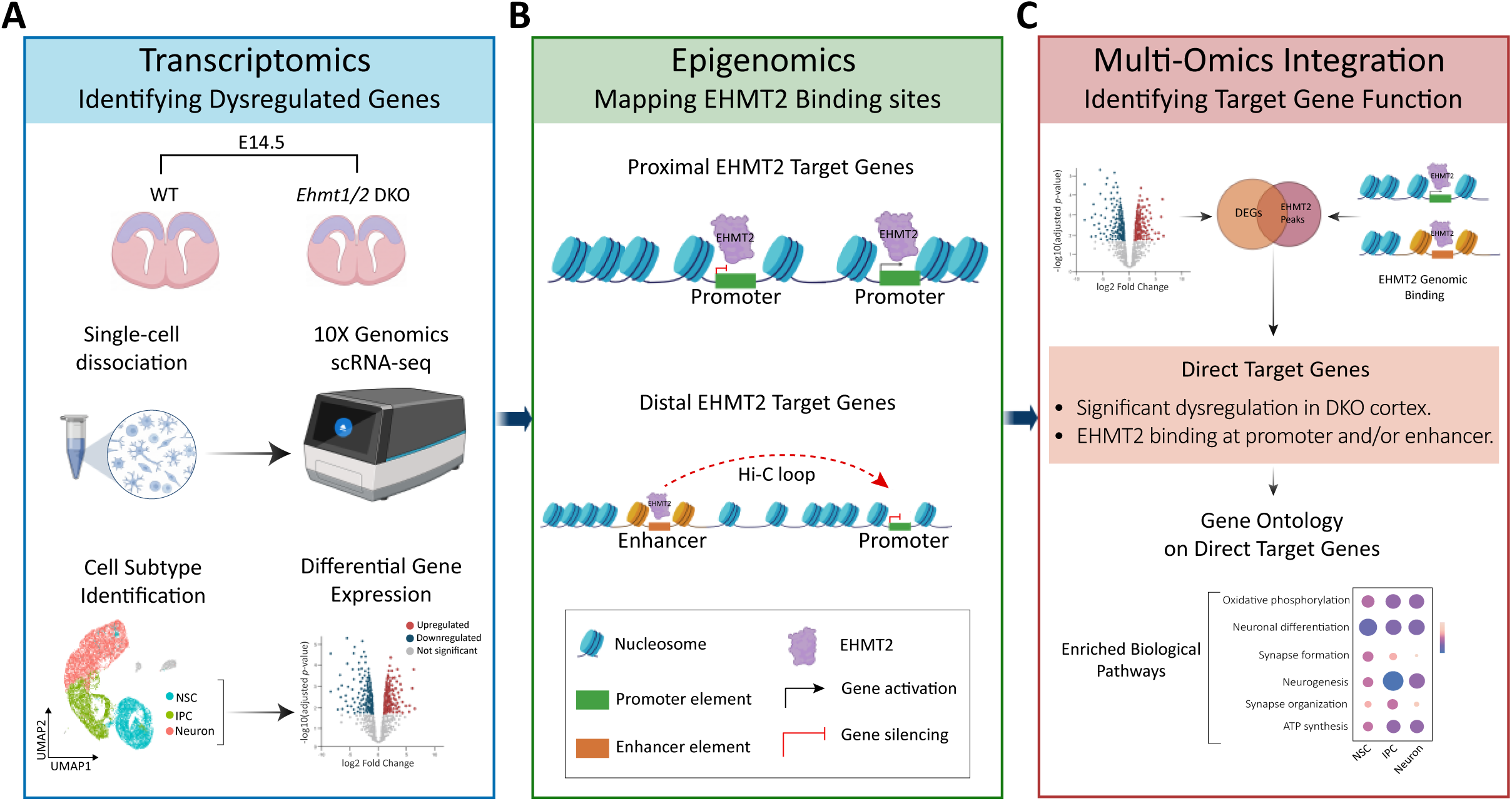
Workflow for identifying putative EHMT2 transcriptional targets in the E14.5 cortex. **(A)** scRNA-seq analysis identified differentially expressed genes (DEGs) between WT and DKO NSCs, IPCs, and neurons. **(B)** ChIP-seq analysis identified EHMT2-binding sites in the E14.5 WT cortex. EHMT2 binding within annotated promoters was used to identify proximal target genes. Previously reported enhancer–promoter interactions identified by Hi-C in the developing cortex were used to identify distal target genes associated with EHMT2-bound enhancers. **(C)** Integrating putative EHMT2 target genes with DEGs between WT and DKO cortices identified genes directly regulated by EHMT2. Gene Ontology analysis was subsequently performed to identify biological processes associated with putative EHMT2 target genes across NSCs, IPCs, and cortical neurons.

**Figure S6.**
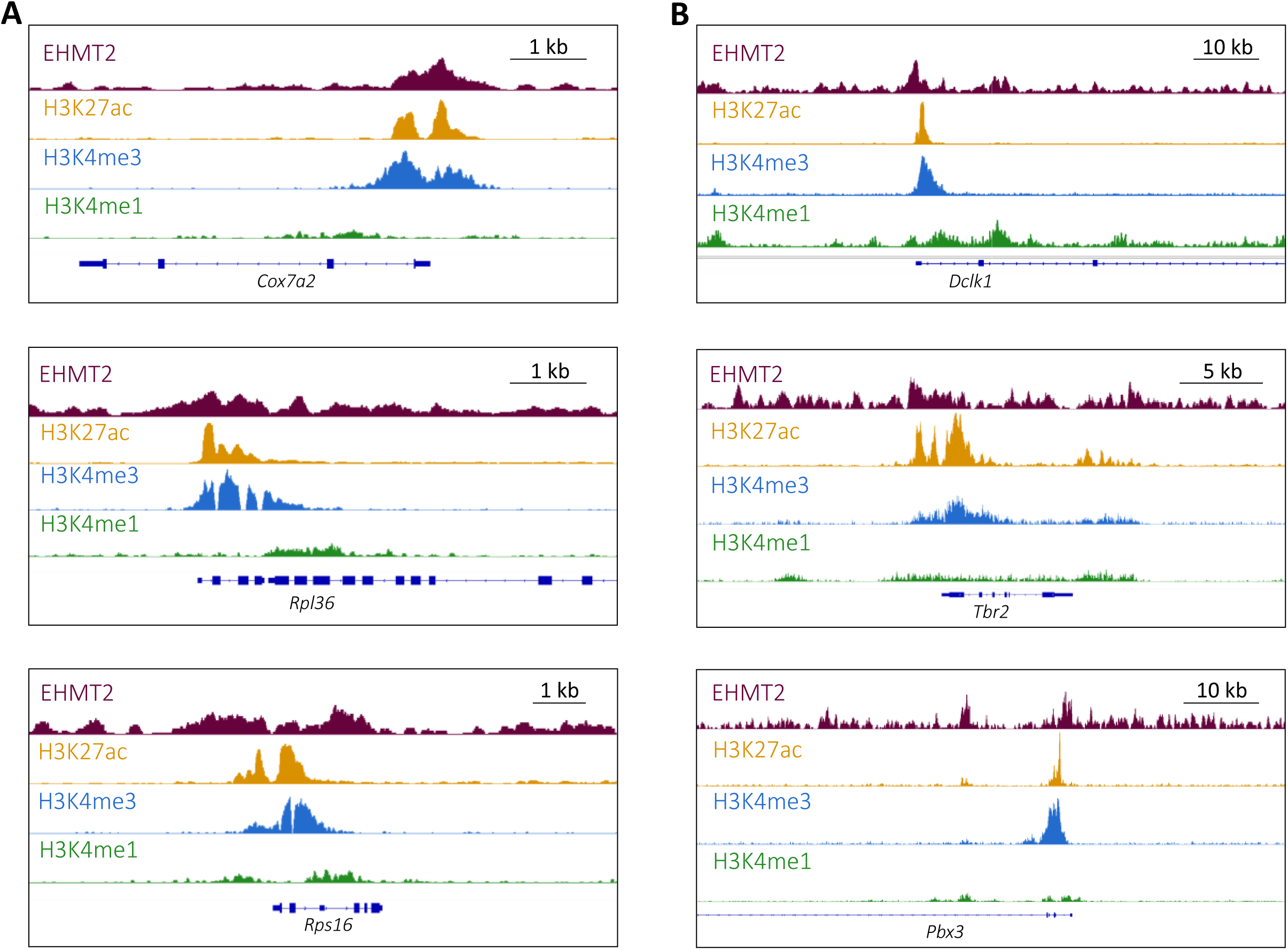
(**A**) Representative genomic tracks showing EHMT2 binding at the promoters of genes associated with oxidative phosphorylation (*Cox7a2*) and ribosome biogenesis (*Rpl36* and *Rps16*). **(B)** Representative genomic tracks showing EHMT2 binding at the promoters of genes associated with neurogenesis (*Dclk1* and *Tbr2*) and upper-layer neuron specification (*Pbx3*). These promoter regions exhibit high H3K27ac and H3K4me3 and relatively low H3K4me1 enrichment.

**Figure S7.**
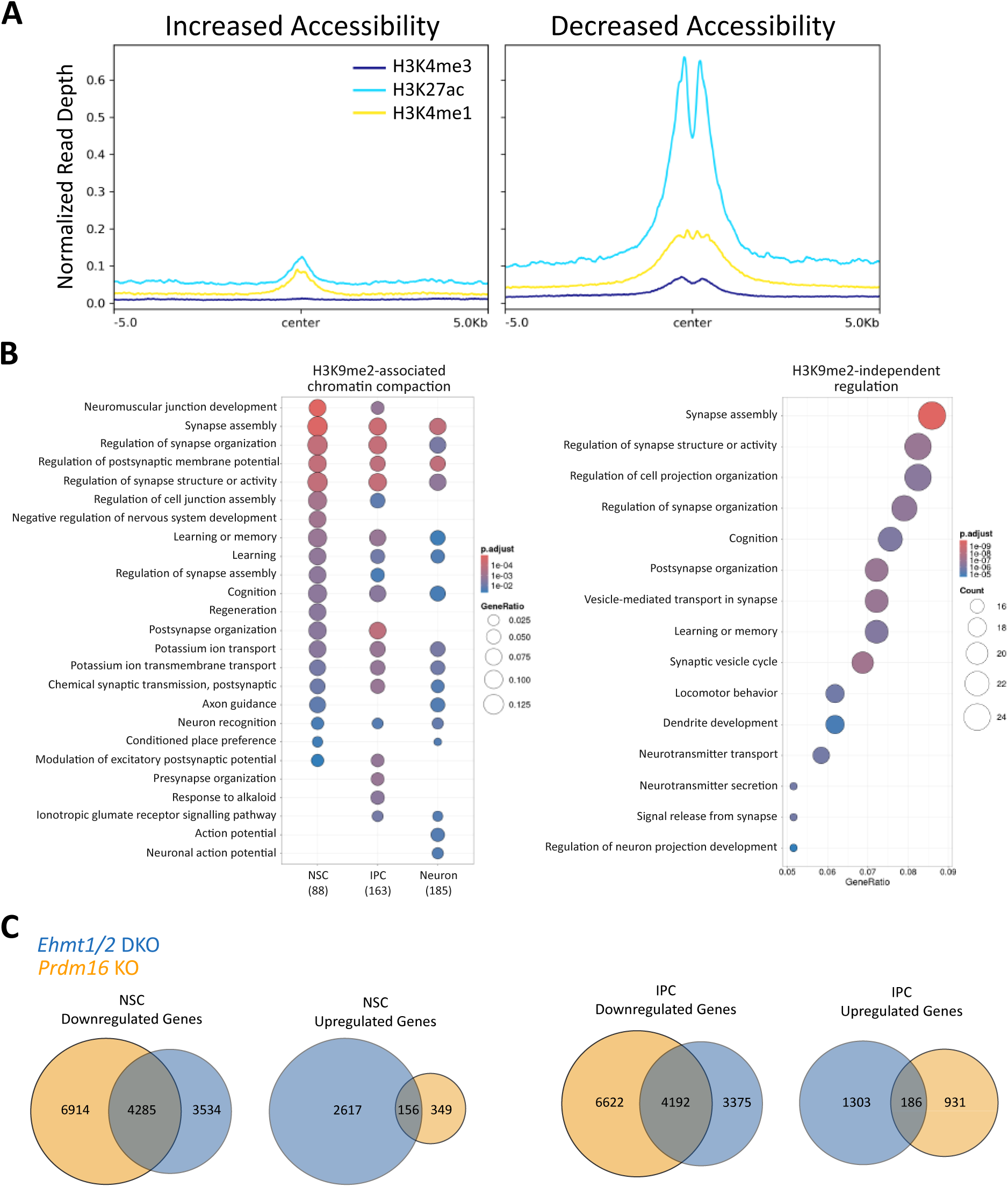
(**A**) Projection profiles showing relatively low levels of the active histone marks H3K4me3, H3K27ac, and H3K4me1 in E14.5 WT cortices at genomic regions that gained accessibility in the DKO cortex (left). Conversely, regions that lost accessibility in the DKO cortex exhibited higher levels of these active histone marks in WT cortices (right). **(B)** Gene ontology (GO) analysis showing enrichment of synapse-related biological processes among genes upregulated and associated with increased chromatin accessibility (left) and among genes upregulated and linked to cluster 2.2, which exhibits minimal to undetectable H3K9me2 enrichment (right). Numbers in parentheses below the left dot plot indicate the total number of genes in each cell type that mapped to significantly enriched GO terms. **(C)** Venn diagrams showing the overlap between downregulated and upregulated genes in NSCs and IPCs from *Ehmt1/2* DKO and *Prdm16* KO cortices.

